# Gene co-expression networks reveal differential developmental modularity in Mammalian limbs

**DOI:** 10.64898/2026.05.07.723669

**Authors:** Aidan O. Howenstine, Karen E. Sears

## Abstract

Mammalian limb development is a complex system involving several signaling centers and coordinated cell behaviors to sculpt a functioning limb capable of the diverse locomotory strategies that mammals exhibit. To investigate the changes in development that facilitate the generation of the wide array of limb phenotypes across mammals, we take a correlation network approach to investigate the developing limbs of mice, bats, and opossums, which represent typical limb development, a novel limb phenotype, and a shift in developmental timing, respectively. Using transcriptomic data of early limb development across these taxa, we build module correlation networks and identify a difference in network connectivity and the distribution of limb development genes across bat limb development. We identify a unique signature of increased modularity in the bat forelimb that is not detected in mouse or opossum. This modularity is not associated with increased specialization of limb development modules, but rather is marked by target limb development genes being spread evenly across several modules. The opossum, with its standard phenotype but altered developmental timing, does not show a difference in modularity relative to mouse. This work points toward the benefit of a network-minded approach to transcriptomic networks, which reveals developmental modularity and potential gene targets for exploration of developmental system evolution.

## Introduction

The development of the mammalian limb has diverged extensively across taxa and between the fore- and hind limbs within individuals, generating a wide range of locomotor strategies. Across mammals, evolutionary changes in limb development have included reductions in digit number, elongation of limb elements, alterations in forelimb-to-hind limb proportions, shifts in developmental timing, and numerous other modifications that together facilitated the remarkable locomotor diversity observed in the clade.

Two well-studied examples of such developmental divergence are found in bats and opossums. In bats, dramatic elongation of the forelimb relative to the hind limb supports the formation of the enlarged hand-wing required for powered flight. In opossums, forelimb and hind limb development are temporally decoupled to accommodate the developmental constraint imposed by precocial birth, enabling newborns to crawl to the mother’s teat immediately after birth.

The developmental changes that underlie these unique phenotypes and strategies have often been studied through the lens of single genetic changes associated with novel traits (Sears et al., 2006; Weatherbee et al., 2006; Doroba & Sears, 2010; Booker et al., 2016; Eckalbar et al., 2016). However, it is increasingly recognized that regulatory divergence and alterations in the wiring of gene interactions also play central roles in the evolution of limb development (Cretekos et al., 2008; Hockman et al., 2008; Cooper & Sears, 2013; Sears et al., 2015). Framing limb development as an integrated developmental process and examining the molecular interaction networks that generate distinct limb phenotypes provides a systems-level perspective for understanding how novel morphologies evolve. This type of gene-network approach to limb development offers a powerful framework for investigating how the architecture of a conserved developmental program can be modified to produce diverse phenotypes, such as the elongated forelimb of bats or the heterochronic shifts in fore- and hind limb development observed in opossums.

### Network Investigation of Limb Development

Comparing how developmental gene networks are organized across taxa with divergent limb morphologies requires a method tractable across non-model species at matched developmental stages. Gene co-expression networks (CNs), built directly from transcriptomic correlations, offer exactly this – a comparative view of network architecture without requiring the extensive regulatory interaction mapping that a full gene regulatory network (GRN) demands. Recent single-cell studies have begun to resolve pieces of this system by characterizing the cell populations underlying bat wing formation, resolving the cellular and genetic basis of developmental heterochrony in the opossum, and establishing cross-species homology in a spatial comparison of mouse and human limb development (Zhang et al., 2023; Lyu et al., 2025; Menchero et al., 2025; Schindler et al., 2025). However, no existing dataset at this resolution offers a stage-matched time series comparing multiple divergent limb types across both fore- and hind limb.

CNs are well suited to this comparative scale. Relying only on bulk transcriptomic data makes cross-species comparison tractable while providing a topological view of developmental programs that can motivate more targeted regulatory investigation and inform theories of how developmental networks evolve. This approach identifies candidates whose co-expression patterns may be associated with species-specific morphological variability and allows us to investigate how network connectivity across taxa with extremely different developmental strategies may be restructured to achieve phenotypic divergence.

We generate and compare correlation networks of bat (*Carollia perspicillata*), opossum (*Monodelphis domestica*), and mouse (*Mus musculus*). The specific aspects of limb development we focus on pattern and sculpt the developing limb: initiation of the limb bud; the Apical Ectodermal Ridge (AER), which establishes proximal-distal (PD) identity; the Zone of Polarizing Activity (ZPA), which establishes the anterior-posterior (AP) axis of the limb; outgrowth and extension of limb tissue; chondrogenesis; and apoptosis of interdigital tissues (Kronenberg, 2003; Fernandez-Teran & Ros, 2008; Towers & Tickle, 2009; Zeller et al., 2009; Hernández-Martínez & Covarrubias, 2011; Duboc & Logan, 2011; Long & Ornitz, 2013; Petit et al., 2017; Tickle & Towers, 2017).

For this study, the bat provides an example of morphological divergence between the forelimb (FL) and hind limb (HL), with its greatly elongated long bones along the PD axis, widened hand along the AP axis, expanded AER and ZPA domains, and unique retention of interdigit tissues to form the wing (Sears et al., 2006; Weatherbee et al., 2006; Cretekos et al., 2008; Hockman et al., 2008). The opossum provides an example of limb heterochrony, with a forelimb that develops far in advance of the hind limb to accommodate its early birth and necessary crawl to its mother’s teat to survive (Lillegraven, 1975; Morris et al., 2024). The opossum also exhibits a reduced physical AER in the forelimb, though its function as a signaling center for limb outgrowth remains (Doroba & Sears, 2010).

We leverage the mouse model to compare these non-model taxa with a baseline expectation of what a more ’typical’ mammalian limb phenotype and developmental timing may look like. We use stage-matched, bulk RNA-seq data across mouse, bat, and opossum to investigate how the architecture of co-expression networks changes across evolutionary divergent limb morphologies and extract candidate genes for further investigation in a regulatory context.

## Methods

### Target gene identification

To inform our expectations of network connectivity and which genes we may expect to vary between taxa and between limbs, we conducted an exhaustive literature search of genes identified in early limb development. We searched PubMed (pubmed.ncbi.nlm.nih.gov/) and Google Scholar (scholar.google.com) for peer-reviewed articles discussing genes involved in limb bud initiation and paddle formation. Searches for the final analyses took place between December 2025 and April 2026. Target categories included limb initiation, formation of the AER, establishment of the AP axis, establishment of the PD axis, chondrogenesis, limb outgrowth, and interdigit cell death. We targeted these processes because we expect them to differ most among bat, mouse, and opossum (Cooper & Sears, 2013; Doroba & Sears, 2010). Results from this search are reported in Table S1 and are used to filter, reduce, and investigate our module network results. We refer to these 165 curated genes as target genes throughout, denoting genes with established roles in limb development rather than known regulatory targets of a specific factor.

### RNA-sequence data processing

Data were obtained from GSE71390, first published by Maier et al., 2017. Of these data, we analyze 3 species –*Mus musculus*, *Monodelphis domestica*, and *Carollia perspicillata,* which we refer to as mouse, opossum, and bat respectively. The fore- and hind limb of each species has been sampled across 3 time points – the ridge, bud, and paddle stages of limb development. These stages cover the very beginning of limb development: the ridge stage marks the initial outgrowth of mesoderm from the flank, the bud stage begins outgrowth of this ridge, establishing the PD and AP axes, and the paddle stage marks the beginning of chondrogenesis of the long bones and the formation of the digits (Maier et al., 2017; Wanek et al., 1989). We use reference genomes GRCm39 (UCSC, accessed February 2024), MonDom1 (UCSC, accessed February 2024), and mCarPer1.2 (Bat1K Consortium/Senckenberg Research Institute, accessed February 2024), for mouse, opossum, and bat samples, respectively. Bat gene annotations were obtained from a TOGA-based annotation for the mCarPer1.2 assembly (Kirilenko et al., 2023, Senckenberg Genome Browser, accessed March 2024), using the human (GRCh38/hg38) reference. Of 18,773 annotated genes, 16,507 (87.9%) were classified by TOGA as strict one-to-one human orthologs; the remainder reflect one-to-many, many-to-one, or many-to-many relationships (n = 1,482; 7.9%) or the absence of an identifiable bat ortholog (n = 589; 3.1%), or were not classified by TOGA (n = 195; 1.0%). Independently, all 18,773 annotated genes were queried against Ensembl BioMart to assign a corresponding mouse gene homolog for cross-species comparison; 15,586 (83.0%) received a mouse homolog assignment regardless of TOGA orthology class, while the remaining 3,187 retained their original bat/human identifier. None of the 165 curated limb-development target genes (Table S1) are affected by non-one-to-one orthology classifications (Table S1a).

Sequences were inspected using fastqc. Reads were trimmed using Trimmomatic v 0.39 (Bolger et al., 2014). Genome alignment was performed using STAR v2.7.9a (Dobin et al., 2013). Aligned reads were indexed with Samtools, and read counts generated using HTSeq v3.7.2 (Anders et al., 2015).

All gene reads were homologized to mouse genes to allow cross-taxa comparisons – this necessarily removes some key genes from the opossum and bat due to the increased quality of annotation for the mouse genome (Jin et al., 2019; Miller et al., 2010; Mueller et al., 2017). This approach is supported by recent single-cell evidence of substantial developmental homology between mouse and human limb development, lending confidence that mouse-referenced homology is a reasonable basis for cross-species comparison in these divergent taxa (Zhang et al., 2023). This permits the direct comparison of correlations and connectivity of our three species on network architecture generated across all species – ensuring we are capturing similarities in limb development across our samples first, before assessing variation.

### Co-expression Network Analyses

Network analyses were performed using the package WGCNA in R (Langfelder & Horvath, 2008). Raw counts were log transformed, and the goodSampleGenesMS function was used for further quality control of our reads. After filtering for homologs and cleaning with goodSampleGenesMS, we end up with 10,562 genes for FL and 10,554 genes for HL.

### Quality control

We check sample clustering and remove poorly clustering samples, leaving us with n = 7 for mouse, n= 11 for opossum, and n = 7 for bat for our forelimb data, and n = 7 each for mouse, opossum, and bat hind limb data. Sample breakdown and filtering is included in Table S2.

### Network generation

Figure 1 provides a conceptual overview of this network-generation and comparison workflow. We constructed consensus networks for FL and HL samples using the blockwiseConsensusModules function of WGCNA, alongside independent species-specific networks constructed using blockwiseModules (Table S3)(Langfelder & Horvath, 2007). The consensus networks identify modules of genes whose co-expression patterns are shared across all 3 taxa, providing a cross-species reference for module composition (Figure 2) and identifying target genes that lack consistent module assignment across taxa (Table 3). Consensus network construction by necessity restricts analysis to the gene set present in all 3 taxa, leaving a more limited yet conserved set of genes to compare. The species-specific networks built from this set identify modules within each taxon independently and are used for the comparative analyses described below.

**Figure 1:**
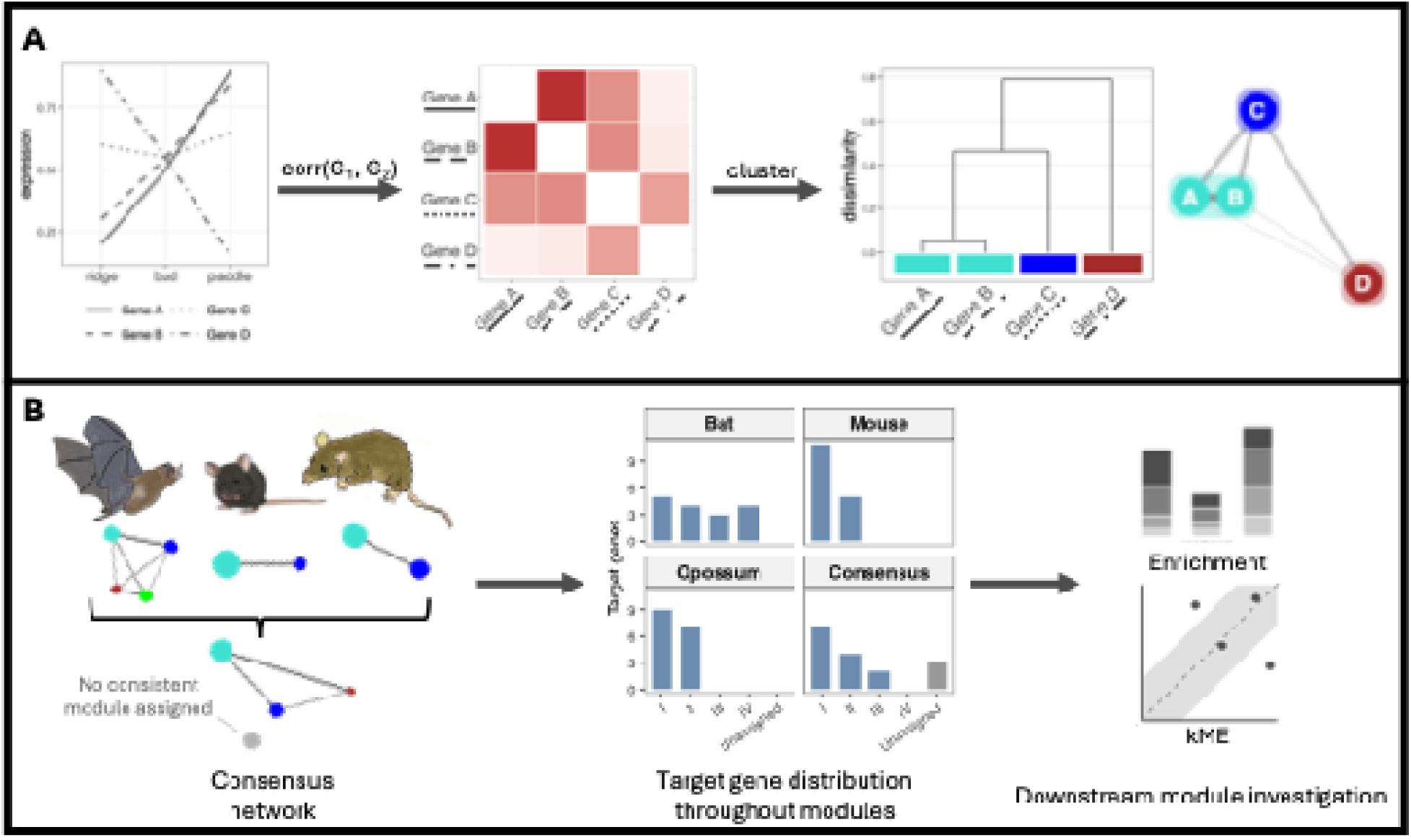
Conceptual workflow for network construction and cross-species comparison. (A) How two genes end up in the same module, illustrated with a hypothetical four-gene example (not real data). Gene expression is compared pairwise across developmental stages via correlation, genes are hierarchically clustered by dissimilarity, and clusters are assigned module identity; genes with correlated trajectories (Gene A, Gene B) form a shared module, while an uncorrelated gene (Gene C) and an anti-correlated gene (Gene D) are each assigned to their own separate module. (B) Networks are built independently for each species from that species’ own within-limb expression trajectories, following the process shown in (A); cross-species comparison happens only afterward with a fourth, pooled consensus network built from the gene set shared across all three species – genes are never grouped by cross-species expression level or by forelimb – hind limb similarity during network construction itself. Unassigned genes are excluded from consensus modules (see Table 3). Module composition for each species-specific and consensus network is summarized (see Figure 2). Enrichment and kME analyses are then conducted on the modules of each species-specific and consensus network (see Figures 3 – 5). Downstream analyses investigate patterns within these modules. Modules and colors in this figure are illustrative and do not correspond to module colors used elsewhere in the manuscript.

**Figure 2:**
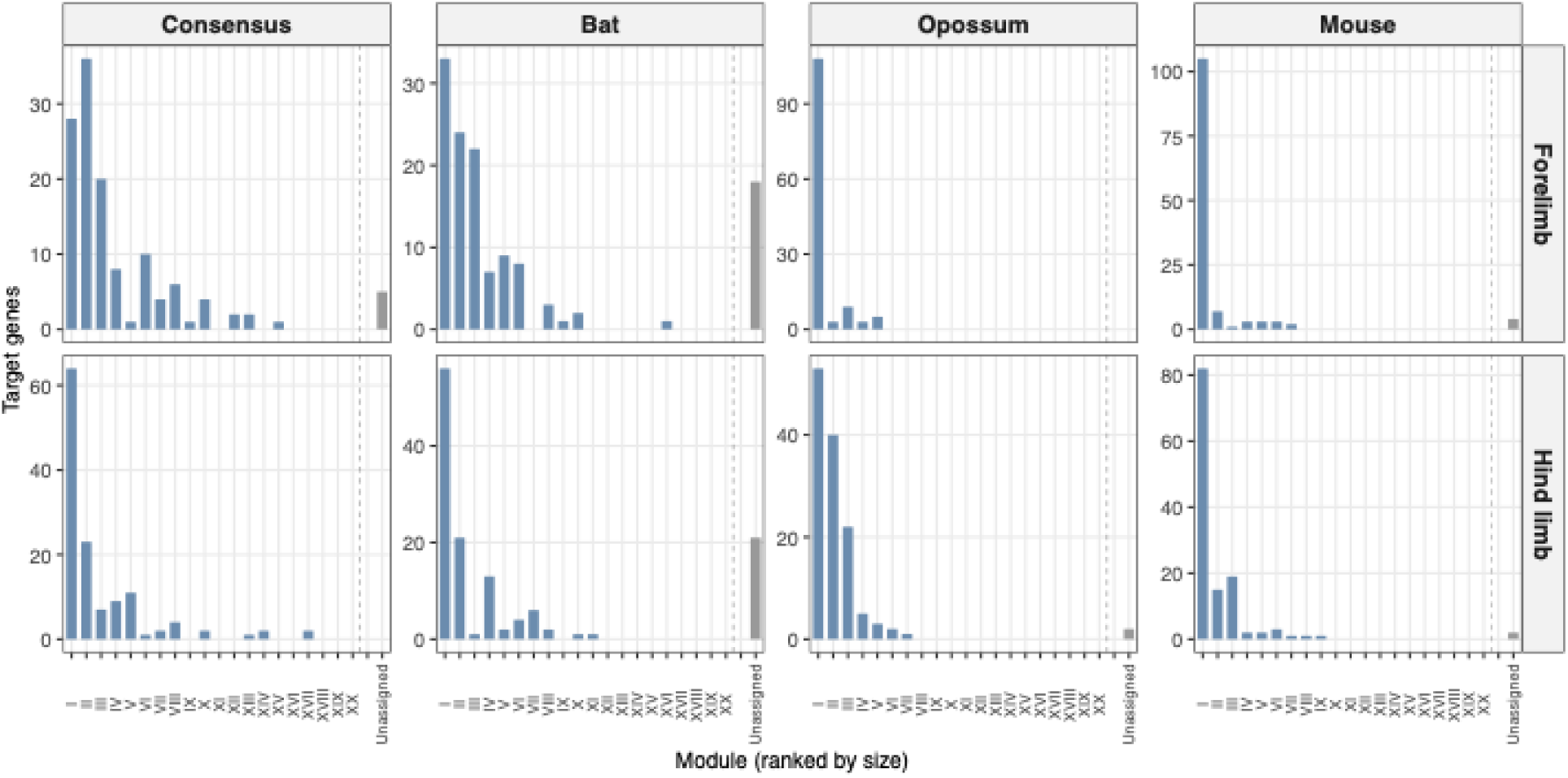
Target gene distribution across co-expression modules. Bar height shows the number of curated limb-development target genes assigned to each module, faceted by network (Consensus, Bat, Opossum, Mouse) and limb (Forelimb, Hind limb). Modules are ordered by rank in total network size (module I = largest module by overall gene membership in that network), not by target-gene count – target-gene distribution does not necessarily track overall module size, and divergence between the two is itself informative (see Results, whole-network vs. target-gene comparison). The final bar (Unassigned), separated from ranked modules by a dashed line, shows target genes that do not cluster with any detected module in that network (see Table 3). Y-axes are scaled independently per panel.

The pickSoftThreshold function indicates imperfect fits for all of our taxa, particularly mouse and opossum, but we use a soft power threshold (a parameter controlling how strongly gene-pair correlations are weighted) of 12 in line with WGCNA developer recommendations for comparative datasets where individuals do not all reach the typical scale-free fit threshold, balancing scale-free fit with preservation of connectivity across taxa. All networks were constructed as signed to distinguish positive from negative correlation within modules – this prevents highly negative correlations from clustering together, so we identify only modules of genes correlated in the same direction (Mason et al., 2009). We set minModuleSize to 40, and mergeCutHeight to 0.30, to prevent small modules from overcrowding our analysis. For species-specific networks, we additionally used corType = “bicor” and maxPOutliers = 0.1 for robustness to expression outliers, as our sample size is at the low end of WGCNA’s capacity and benefits from a more conservative approach (Langfelder & Horvath, 2012). Module color labels are assigned independently within each network and do not indicate cross-network correspondence. Network dendrograms are provided in supplementary figures 1-4.

### Enrichment analysis

We perform gene ontology (GO) enrichment on our modules using the R packages clusterProfiler and org.Mm.eg.db (Wu et al., 2021). We filter our results for target GO terms involving limb development. We generate custom functional categories based on an exhaustive literature search of limb-associated genes, and test for enrichment of these targets. Many genes function in multiple developmental processes and are accordingly assigned to multiple categories (65 of 156 genes are assigned to >1 category, for an average of 1.56 categories per gene). Categories were thus not treated as wholly independent in interpretation. We filter these results for modules that have more than 5 genes present for each category.

With our literature-sourced limb development gene categories (Table S1), we perform dispersion tests of module-category combinations based on Shannon entropy to examine the effective number of modules each developmental category is spread between (Tables 1, S6, and S7).

### Module connectivity

To identify hub genes for each module, we calculate kME (module eigengene connectivity), a measure of how strongly each gene’s expression tracks its assigned module’s overall expression trend. Each module’s overall trend is summarized by its module eigengene, a single representative expression profile for all genes in that module; kME is the Pearson correlation between a gene and this eigengene. High kME indicates greater importance and correlation within a module. We filter these results to identify the top 10% kME genes within each module, and define these genes as hub genes, which are those that are most strongly correlated with their assigned module. We investigate these results to identify genes that are unique hubs within their species and limbs and highlight those that gain hub status in opossum or bat relative to mouse.

To identify uniquely high kME values between limbs and taxa, we plot FL vs HL kME for each taxon, as well as bat vs mouse, bat vs opossum, and opossum vs mouse kME for both FL and HL to identify differential kME in our samples. We use a threshold of 0.3 to mark notable difference in kME across networks (Horvath & Dong, 2008; Langfelder & Horvath, 2008).

## Results

### Module gene distribution

Across the six species-limb networks, target limb genes (Table S1) align into several modules; we focus on the largest, numbered I – VI by decreasing size within each network.

Limb gene composition across forelimb and hind limb networks for bat, opossum, and mouse reveal that bat limb development is characterized by a more even distribution of limb genes across developmental modules (Figure 2). In the bat forelimb, the 3 largest modules – modules I, II, and III – hold 33, 24, and 22 target limb genes, respectively. This is in contrast to the opossum and mouse, where the largest module holds over 100 target limb genes and the next largest modules hold less than 10. The comparison of the networks for the hind limbs of bat, mouse, and opossum shows a slightly more nuanced pattern, with similar distributions of limb genes across the major modules, but again the bat exhibits a more even spread of limb genes across modules (Figure 2).

Bats also show a different pattern of unassigned target genes, which includes those genes that do not tightly cluster with any specific module. Specifically, the bat has far more genes unassigned than any other species. The bat FL and HL have 18 and 21 unassigned target genes, respectively, while the opossum FL and HL have 0 and 2, respectively, and the mouse FL and HL have 4 and 2, respectively (Table 3). This indicates that a substantially greater amount of canonical limb development genes do not cluster as expected in the bat based on our findings in mouse and opossum.

### Module category enrichment

To understand whether the observed difference in the module assignment of limb-associated genes in bats and other examined species is specific to the developmental categories that we have identified, (e.g., AER, AP patterning, chondrogenesis, interdigit apoptosis, outgrowth, and limb initiation), we performed dispersion tests of our categories within modules to examine the effective number of modules each category is spread across. Across dispersion tests of category enrichment (Table 1), GO enrichment (Table S4a, S4b), visual analysis (Figure 3), and Fisher’s-tests (Table 2), we find that limb development categories are distributed evenly across the largest modules in most networks, and there are few distinct module-category pairings.

**Figure 3:**
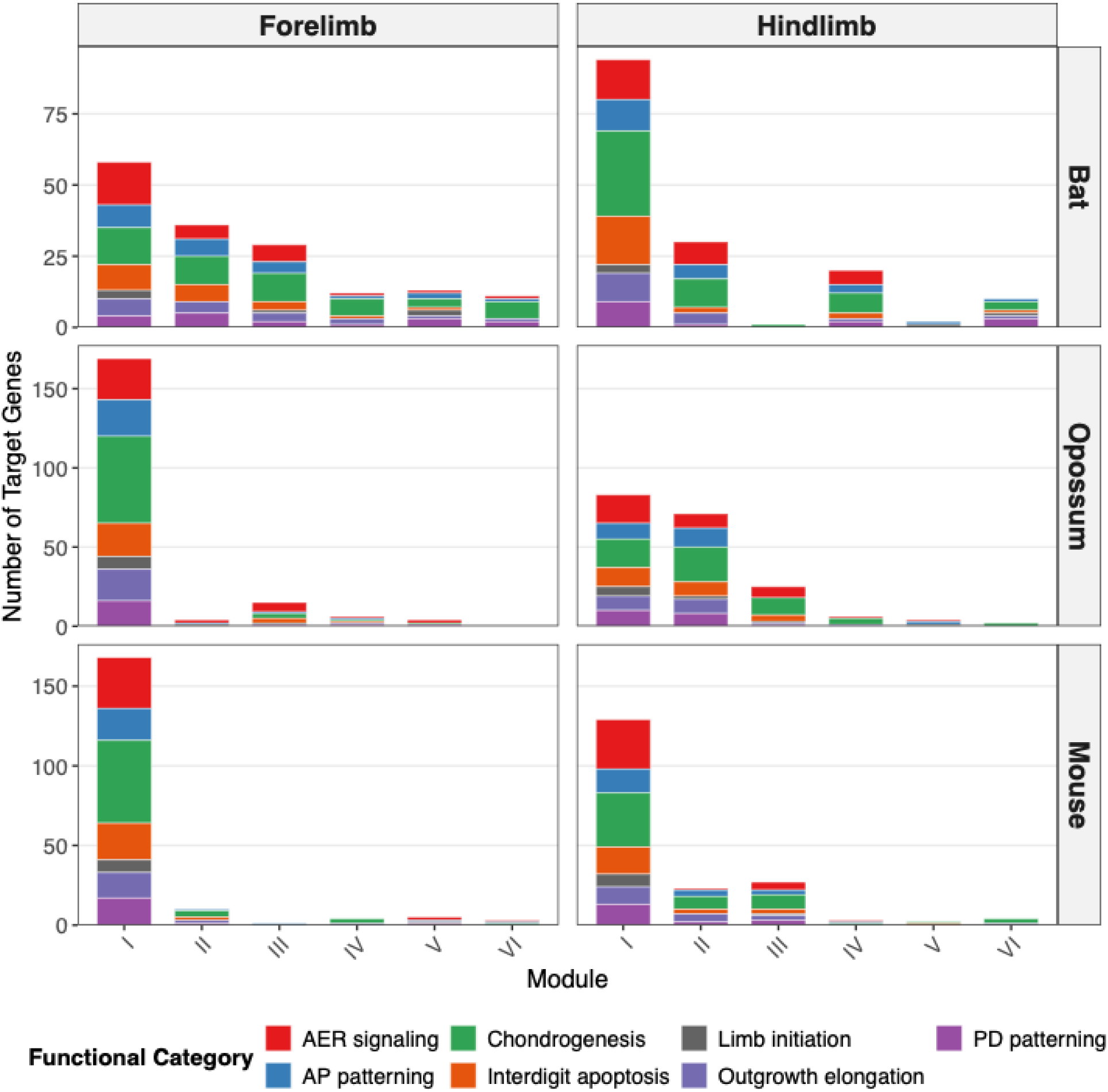
Functional category composition of the largest co-expression modules across species and limb networks. Stacked bars show the number of limb development target genes assigned to modules I - VI, colored by functional category membership. Gene counts reflect functional category membership and are not mutually exclusive, as individual genes may belong to multiple categories.

**Table 1.** Per-network dispersion of limb-development gene categories across modules. Effective number of modules computed as exp(Shannon entropy of category gene counts across assigned modules). Higher values indicate broader distribution. Evenness ranges 0–1, with 1 indicating perfect even spread across occupied modules.

| **Network** | **Categories tested** | **Effective N modules (median)** | **Effective N modules (mean)** | **Median evenness** |
| --- | --- | --- | --- | --- |
| Bat FL | 7 | 4.62 | 4.97 | 0.86 |
| Bat HL | 7 | 3.59 | 3.53 | 0.78 |
| Opossum FL | 7 | 1.70 | 1.76 | 0.44 |
| Opossum HL | 7 | 2.77 | 2.99 | 0.79 |
| Mouse FL | 7 | 1.69 | 1.67 | 0.40 |
| Mouse HL | 7 | 2.68 | 2.61 | 0.64 |

**Table 2.**
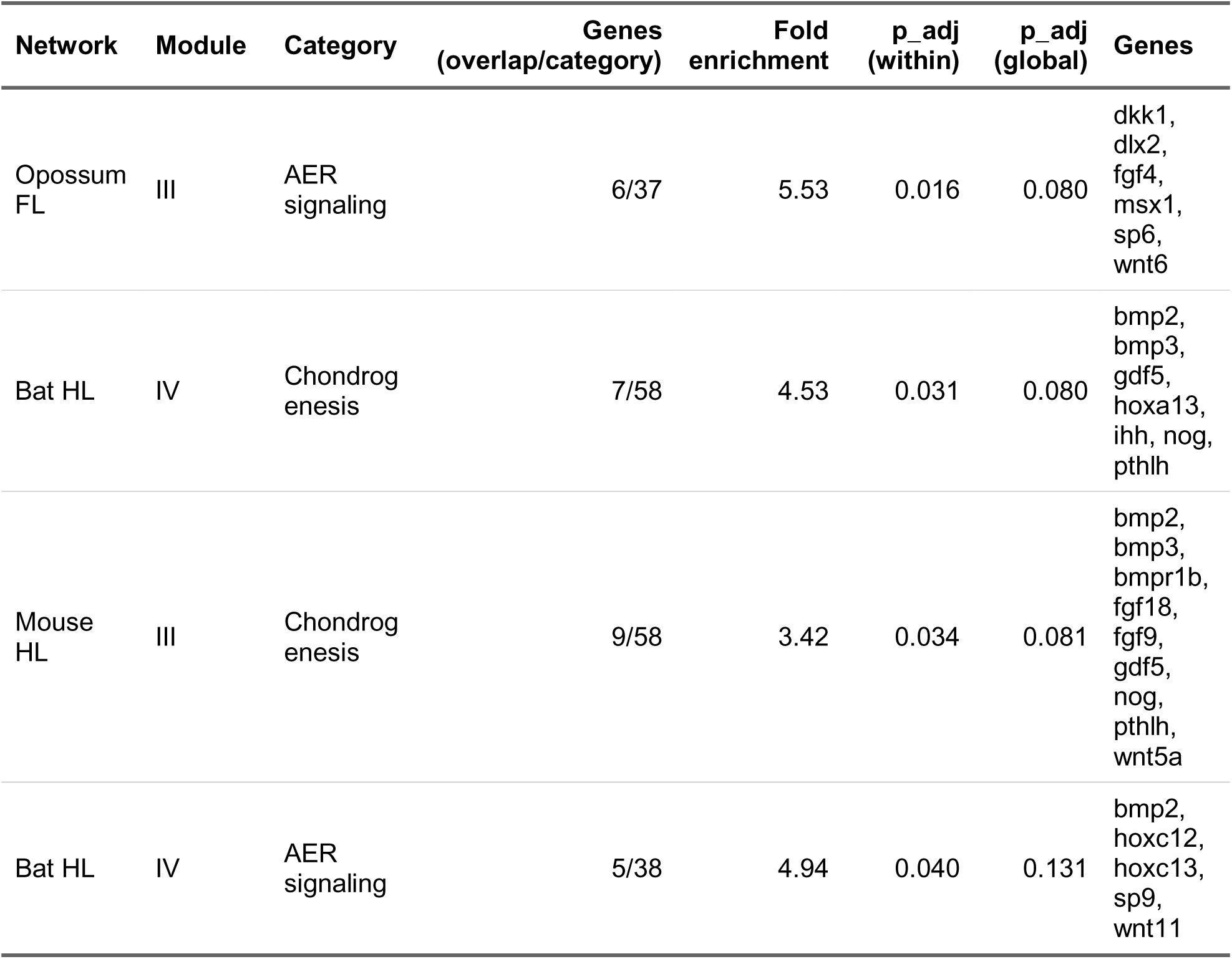
Significant module-category enrichments under Fisher’s exact test. Filters: overlap > 5 genes, fold enrichment > 2, within-network adjusted p-value < 0.05. Of 191 module-category combinations with non-zero gene overlap tested across the 6 species-specific networks, 4 reached significance under within-network FDR; none surpassed the more stringent global FDR threshold. A significant enrichment was also identified in the Consensus network (Table S5); this table is scoped to the species-specific networks and does not include that result.

The module distribution for bat shows a more dispersed, even spread of limb genes and developmental categories across both the forelimb and hind limb relative to the mouse and opossum, which show fewer modules and greater density of developmental categories in the largest module, module-I (Figure 3). Dispersion results show that the bat FL is most evenly dispersed, with an average of 4.97 effective modules per category, followed by the bat hind limb with 3.53 effective modules (Table 1). Both mouse and opossum forelimb have under 2 effective modules per category, and for the hind limb under 3 effective modules per category. Based on these results, the bat forelimb shows more dispersion than its hind limb, while the opposite holds true for mouse and opossum fore- and hind limbs (Table 1). With only 7 categories, the Wilcoxon signed-rank tests are limited in their resolution, but still show significant differences between bat and mouse or opossum networks, as well as between bat FL and HL networks (p = 0.046), but not between mouse and opossum FL (p = 0.834) or HL networks (p = 0.650; Table S8).

The categories driving the greater dispersion in the bat FL network relative to those of the FL of mouse and opossum are chondrogenesis (6.61 effective modules), outgrowth (5.61), AP patterning (4.62), and limb initiation (4.46) (Table S7). These categories are spread between 4 and 6 modules in the bat FL, but only between 1-2 modules in both mouse and opossum FL.

GO enrichment for limb-development specific terms supports our independent observation of functional categories being distributed across multiple modules (Table S4b). Narrowing enriched GO terms only to limb-development associated terms reveals that the bat FL shows the greatest distribution of limb associated GO terms, with 27 terms distributed across four modules (modules I, II, III, and VI) and the unassigned category. This is in comparison to 11 terms identified for mouse and opossum forelimb, all in module-I. For the hind limb, all taxa exhibit intermediate patterns, similar to the category enrichment results. Bat exhibits 25 terms across 3 modules, mouse 29 terms across 5 modules, and opossum 19 terms across 4 modules (Table S4b).

Using Fisher’s exact test to examine any categories that are not evenly distributed across modules reveals only a few categories that are significantly represented within a single module. We test for more than 5 genes within a category being present in a module, with fold enrichment > 2, and a within-taxon adjusted p-value < 0.05. We also test a more stringent global FDR correction, which produces no significant enrichment for any of our species-specific networks. In the bat, we only identify the hind limb module-IV as being significantly represented with chondrogenesis and AER signaling genes (Table 2, full results in Table S5). The mouse shows significant enrichment for chondrogenesis genes within module-III of its hind limb, and the opossum shows significant enrichment of AER signaling genes in module-III of its forelimb. Despite this limited significance under strict correction, the underlying dispersion pattern in bat forelimb is independently corroborated by GO-term enrichment using a non-curated annotation source (Table S4b), indicating this reflects statistical power constraints common across our networks rather than an absence of real biological signal of co-expression network differences.

To further test whether this signal reflects genuine network architecture rather than an artifact of gene curation, we compared target-gene module assignments against whole-network module structure (Table S8a). In bat, the greater dispersion of target genes in the forelimb relative to the hindlimb (4.97 vs. 3.53 effective modules) was mirrored at the whole-network level (16 vs. 11 modules and largest module comprising 50.0% vs. 63.8% of network genes in FL vs HL), indicating this pattern reflects broader network architecture rather than target-gene selection. In opossum, the concentration of target genes in a single dominant forelimb module was similarly present at the whole-network level (5 vs 9 modules and largest module comprising 90.6% vs 41.9% of network genes in FL vs HL). We note the opossum forelimb dataset includes 11 biological replicates versus 7 for the hindlimb (Table S2), though the direction of this effect (larger n associated with lower module resolution) suggests this result is not a simple consequence of sample-size difference. In mouse, by contrast, whole-network module structure was nearly identical between limbs (largest module 80.8% vs. 81.1% of network genes), yet target genes showed a significant redistribution away from the dominant forelimb module in the hindlimb (84.7% vs. 65.1% of target genes in the largest module of FL vs HL, p < 0.001), indicating this pattern is specific to the target gene set rather than inherited from whole-network structure. These redistributed genes concentrate disproportionately in the chondrogenesis- enriched mouse hind limb module-III described above (fold enrichment = 4.19, within-network adjusted p = 0.034; Table S5), though this does not survive global FDR correction (adjusted p = 0.081) and we present it as suggestive rather than conclusive.

This result points toward interesting biological differences across bat, mouse, and opossum that remain to be explored. The bat forelimb develops a novel phenotype relative to its hindlimb, and we suggest that both whole co-expression network and the curated limb-target set network reorganization is a feasible pattern associated with this drastic difference. For example, the retained interdigital tissue and extended chondrogenesis of the bat forelimb each plausibly reflect this rewiring (Weatherbee et al., 2006; Hockman et al., 2008; Eckalbar et al., 2016). The opossum, however, develops morphologically similar phenotypes between its fore- and hind limbs, but the timing of their development is drastically different relative. For this reason, we may expect the curated limb-target set network to be similarly constructed to our model system, the mouse, but the broader co-expression network to show differences between the fore- and hind limbs due to their divergent developmental trajectories; the marsupial forelimb is accelerated and hind limb comparatively delayed to support precocial crawling to the pouch (Keyte & Smith, 2010, 2014). This expectation is consistent with the results in the mouse, which exhibit the same pattern to the opossum in the limb-target network, with the hind limb being more modular than the forelimb, but reveals more homogeneity between the broader all-gene co- expression network. This discrepancy between the limb-target network and all-gene coexpression network of the mouse is unexplained by the analyses here, and may reflect a bias introduced by our curated set of limb genes, which are necessarily based primarily on experiments performed in mice. Why the hind limb of both the mouse and opossum are identified as more modular remains unexplained by our data, but we speculate that this may reflect the divergent regulatory architecture of the fore- and hind limbs, particularly in their inputs to begin limb initiation (Rabinowitz & Vokes, 2012).

### Module Eigengene correlations

We next examined whether these network-level patterns revealed target genes with unique co-expression profiles associated with limb development for each examined species. Comparing differences in intramodular connectivity (kME) between FL and HL within species can identify unique genes that may be associated with differential forelimb and hind limb development in each taxon. We identify 14 shifts in kME > 0.3 in bat, 16 in opossum, and 16 in mouse between FL and HL networks, but note a trend of more high-kME genes in the opossum HL and more high-kME genes in the mouse FL, while genes are evenly distributed between FL and HL for bat (Figure 4). For the bat, we identify *Shh*, *Runx2*, and *Fgf18* as uniquely high kME genes in the FL networks. We find *Trp63* to exhibit uniquely negative kME for the FL network relative to the HL network, indicating a negative correlation with its assigned module in the FL that was not identified in the HL. For opossum, we identify *Grem1*, *Hoxa9*, and *Hoxd9* as exhibiting high HL kME in comparison to the FL network, and *Hoxc5* and *MycN* as uniquely high kME in the FL network. Mouse data shows a strong bias toward higher kME in the FL (87 of 126 target genes; paired Wilcoxon test, p = 3.4 x 10Z Z), with *Wnt10b* exhibiting the greatest difference from the HL network, and we identify *Hand1* and *Fgf4* as having uniquely negative FL kME. The developmental categories these outliers span include AER signaling (*Fgf18*, *Fgf4* and *Wnt10b*), patterning (5’ Hox genes and *Hand1*), and chondrogenesis (*Runx2* and *Grem1*).

**Figure 4:**
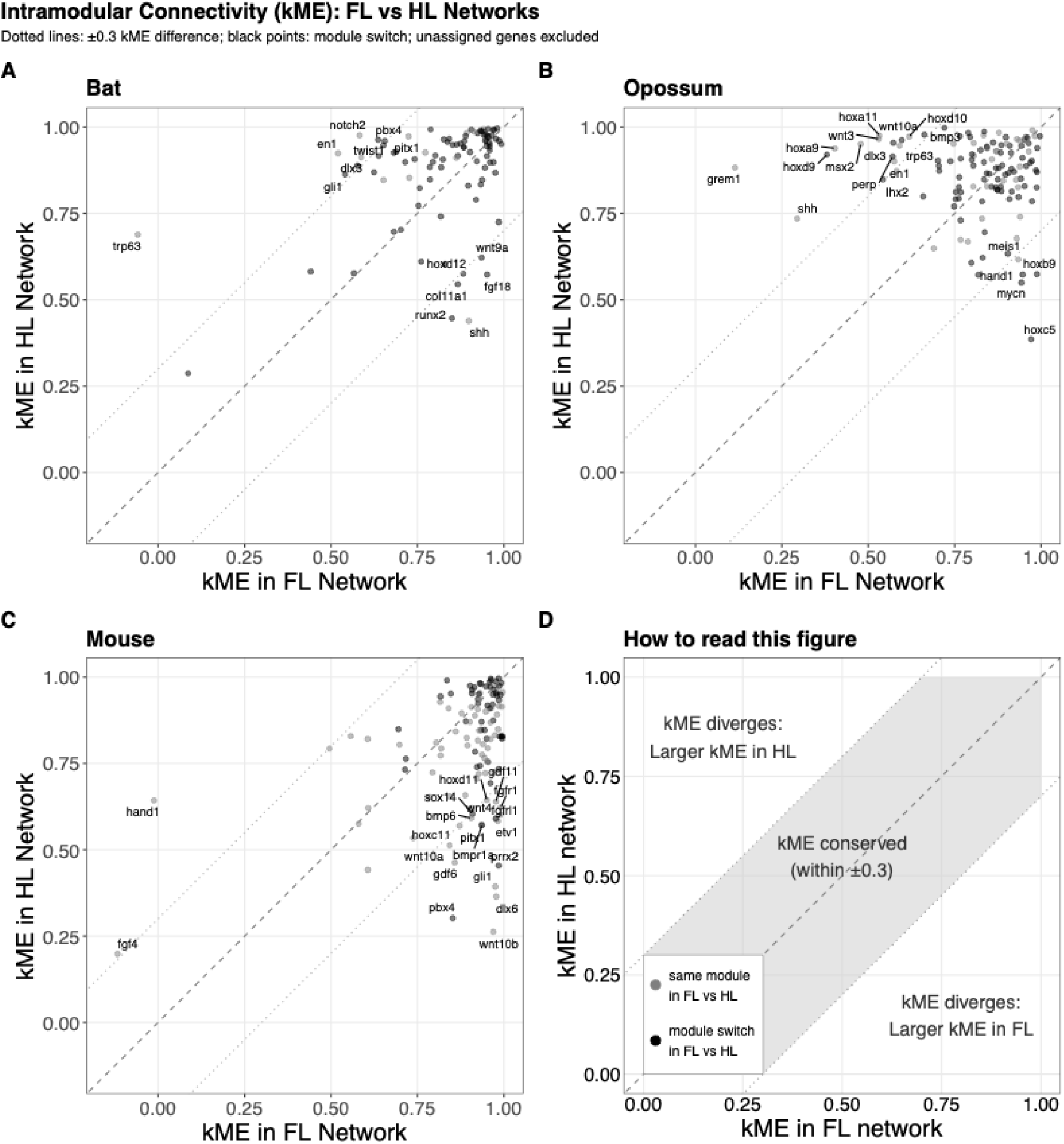
FL vs HL kME across taxa. Each panel plots eigengene correlation (kME) for each gene within its module in the forelimb network against the hind limb network for a single species. (A) Bat, (B) opossum, (C) mouse. Dashed diagonal line indicates equal kME between limb types; dotted lines indicate ±0.3 kME difference threshold. Genes falling outside the dotted lines exhibit limb-specific shifts in module connectivity, and represent strong candidates for characterizing limb-specific reorganization of co-expression networks. Genes with kME below zero on either axis are anti-correlated with their assigned module's eigengene in that network, despite retaining that module assignment. Black points indicate genes that switch module assignment between networks. Labeled genes exceed the 0.3 kME difference threshold. Unassigned genes excluded from analysis. (D) Visual guide to interpreting panels A–C, illustrating the conventions described above.

Comparing kME across taxa can identify genes that may have divergent roles in the coordination of limb development in each species. Relative to mouse and opossum, bat shows very low kME for *Bmp6* in the forelimb, and negative kME for *Trp63*. The only consistently high kME gene for bat relative to mouse and opossum is *Hoxd9*. The opossum FL shows uniquely low *Grem1*, *Hoxa9*, and *Msx2* kME values relative to both mouse and bat. For the hind limb, the bat shows uniquely low kME for *Isl1*, *Bmp6*, and *Runx2*, and high kME for *Dlx6* compared to mouse and opossum. The opossum hind limb shows no genes with >0.3 kME difference from both mouse and bat (Figure 5).

**Figure 5:**
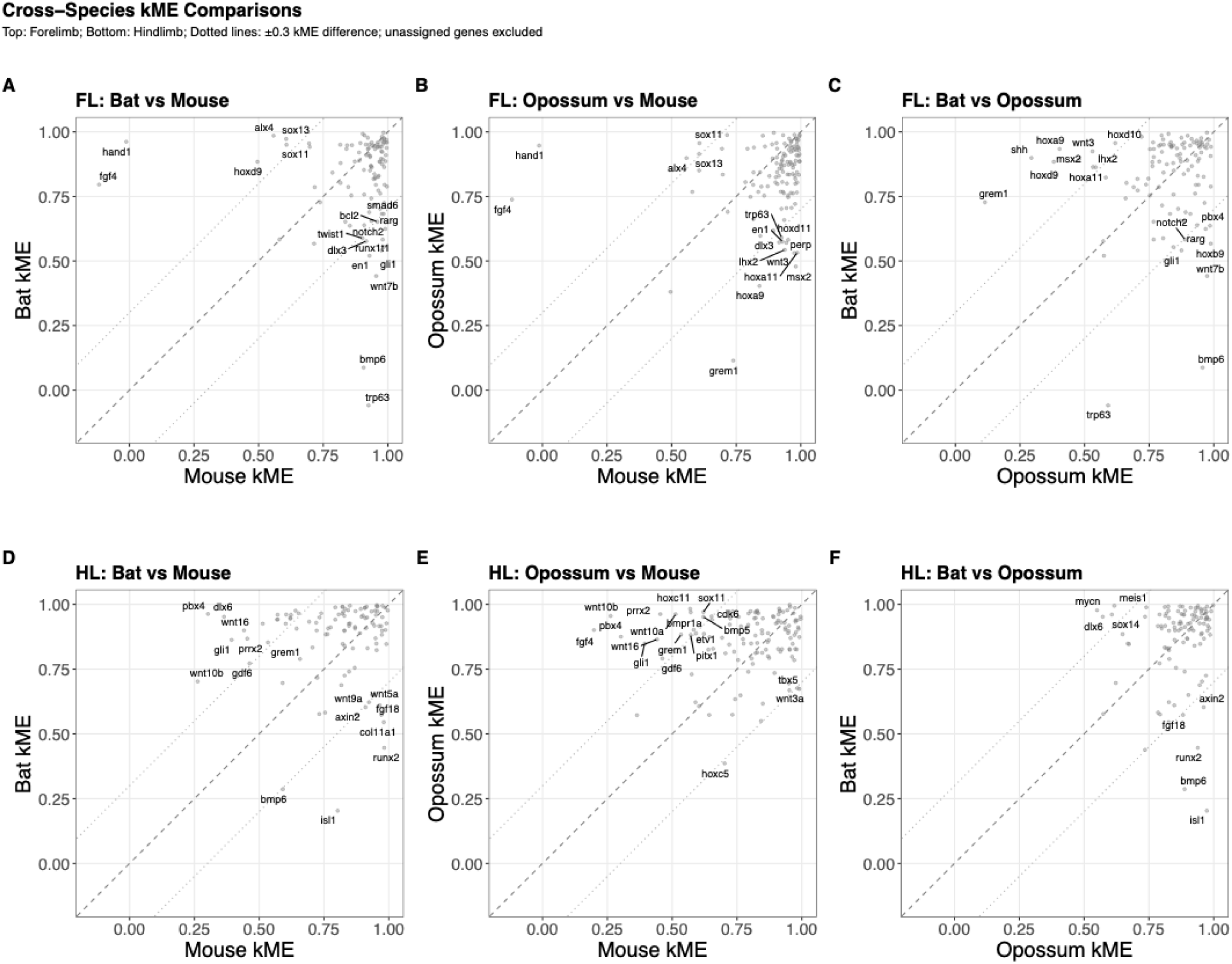
kME comparisons between taxa. Plots gene-module connectivity (kME) values for one species against another within the same limb network. (A-C) Forelimb network comparisons: (A) bat vs mouse, (B) opossum vs mouse, (C) bat vs opossum. (D-F) Hind limb network comparisons: (D) bat vs mouse, (E) opossum vs mouse, (F) bat vs opossum. See Figure 4D for interpretation guide; conventions are identical here, but comparing species instead of FL vs HL. Dashed diagonal line indicates equal kME between species; dotted lines indicate ±0.3 kME difference threshold. Genes falling outside the dotted lines exhibit species-specific shifts in module connectivity, and represent the strongest candidates for characterizing lineage-specific reorganization of limb co-expression networks. Genes with kME below zero on either axis are anti-correlated with their assigned module’s eigengene in that network, despite retaining that module assignment. Labeled genes exceed the 0.3 kME difference threshold. Unassigned genes excluded from analysis.

Several limb development genes fall within the top 10% of kME values for their modules, indicating tight correlations with their assigned modules (Table 4). Based on this, we define these as hub genes. Many of these hub genes differ in bat relative to mouse and opossum. In the bat forelimb, specifically, of note are several posterior Hox genes associated with digit patterning – *Hoxa10, Hoxc10, Hoxd10, and Hoxc13*, and genes involved in the formation of a limb skeleton – *Bmpr1b*, *Gdf5*, *Gdf11*, *Smad4*, *Fgf18*. Of these, *Gdf5* is the only gene that is marked as a hub in both forelimb and hind limb of the bat. Fewer genes are identified as hubs for the opossum, but *Pbx2* and *Isl1* are identified as hub genes in the FL and HL, respectively; both of these genes are involved in coordinating early limb initiation (Capellini et al., 2006; Narkis et al., 2012; Rabinowitz & Vokes, 2012). Hubs that we identify for the mouse include *Tbx5* and Fgf receptor genes in the forelimb, and *Wnt3a* in the hind limb, which are involved in initiation and outgrowth of limb development (Xu et al., 1998; Kawakami et al., 2001; Agarwal et al., 2003; Verheyden et al., 2005; Sheeba & Logan, 2017; Lin & Zhang, 2020).

**Table 3.** Unassigned target limb-development genes in each network. Unassigned genes are those whose expression does not co-vary cohesively with any detected module. Bat networks contain substantially more unassigned genes than opossum or mouse networks.

| Network | Limb | N target genes | Unassigned target genes |
| --- | --- | --- | --- |
| Bat | Forelimb | 18 | bmpr2, dkk1, ets1, fgfr4, fgfrl1, gli2, isl1, jag2, perp, rdh10, smad9, sp9, tbx15, tbx2, twist2, wnt10a, wnt16, wnt5b |
| Bat | Hindlimb | 21 | bmp7, ccne1, dlx1, etv1, fgf16, fgf20, fgf4, glis3, hand1, hhip, hoxc10, hoxc5, hoxc9, lmx1b, pitx2, ptch1, runx1t1, smad9, tbx3, tbx5, twist2 |
| Opossum | Forelimb | 0 |  |
| Opossum | Hindlimb | 2 | dkk1, fgf16 |
| Mouse | Forelimb | 4 | fgf16, hoxc5, irx3, shh |
| Mouse | Hindlimb | 2 | shh, tbx2 |
| Consensus | Forelimb | 5 | dkk1, hand1, hoxc10, wnt4, wnt6 |
| Consensus | Hindlimb | 0 |  |

**Table 4.**
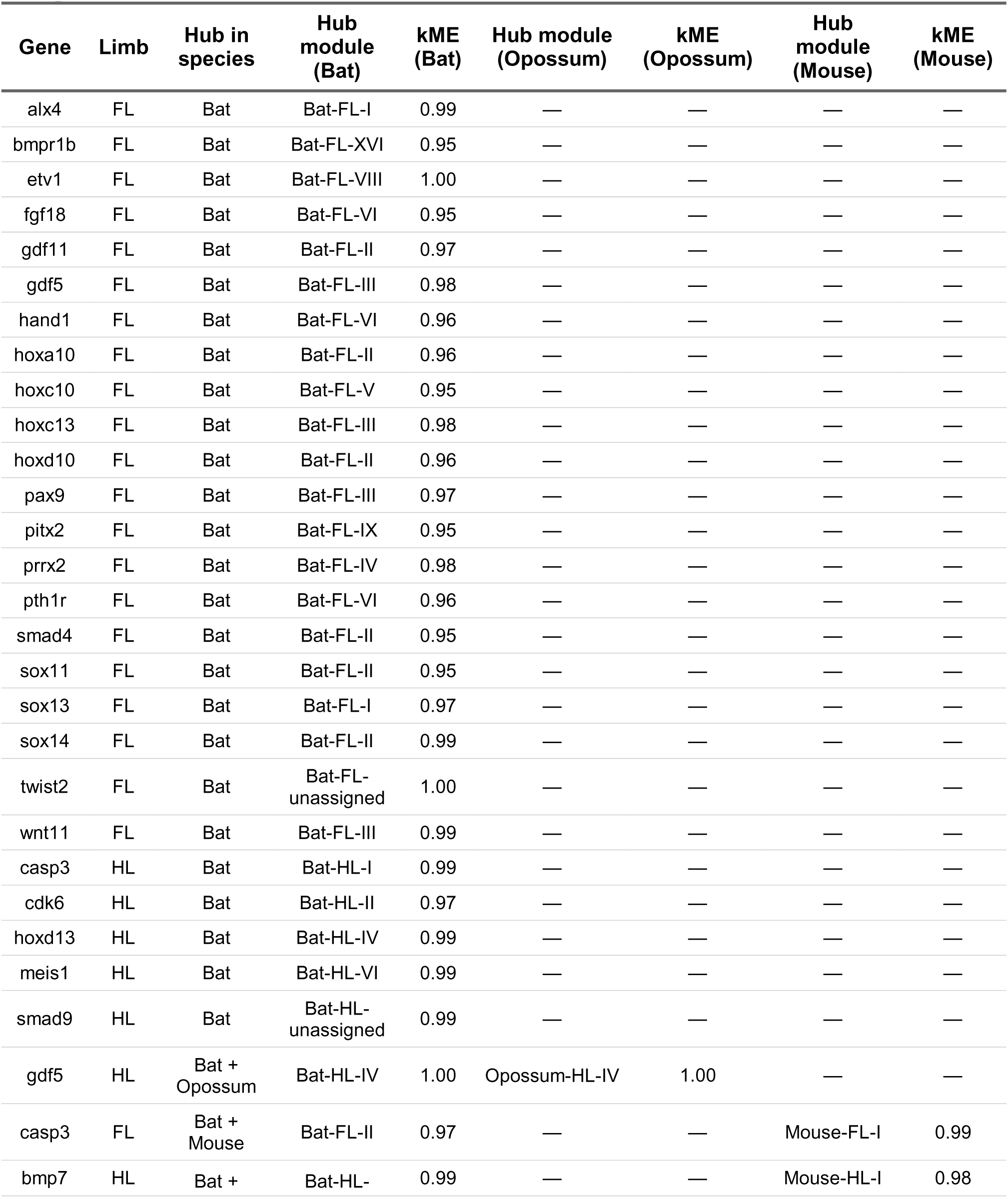
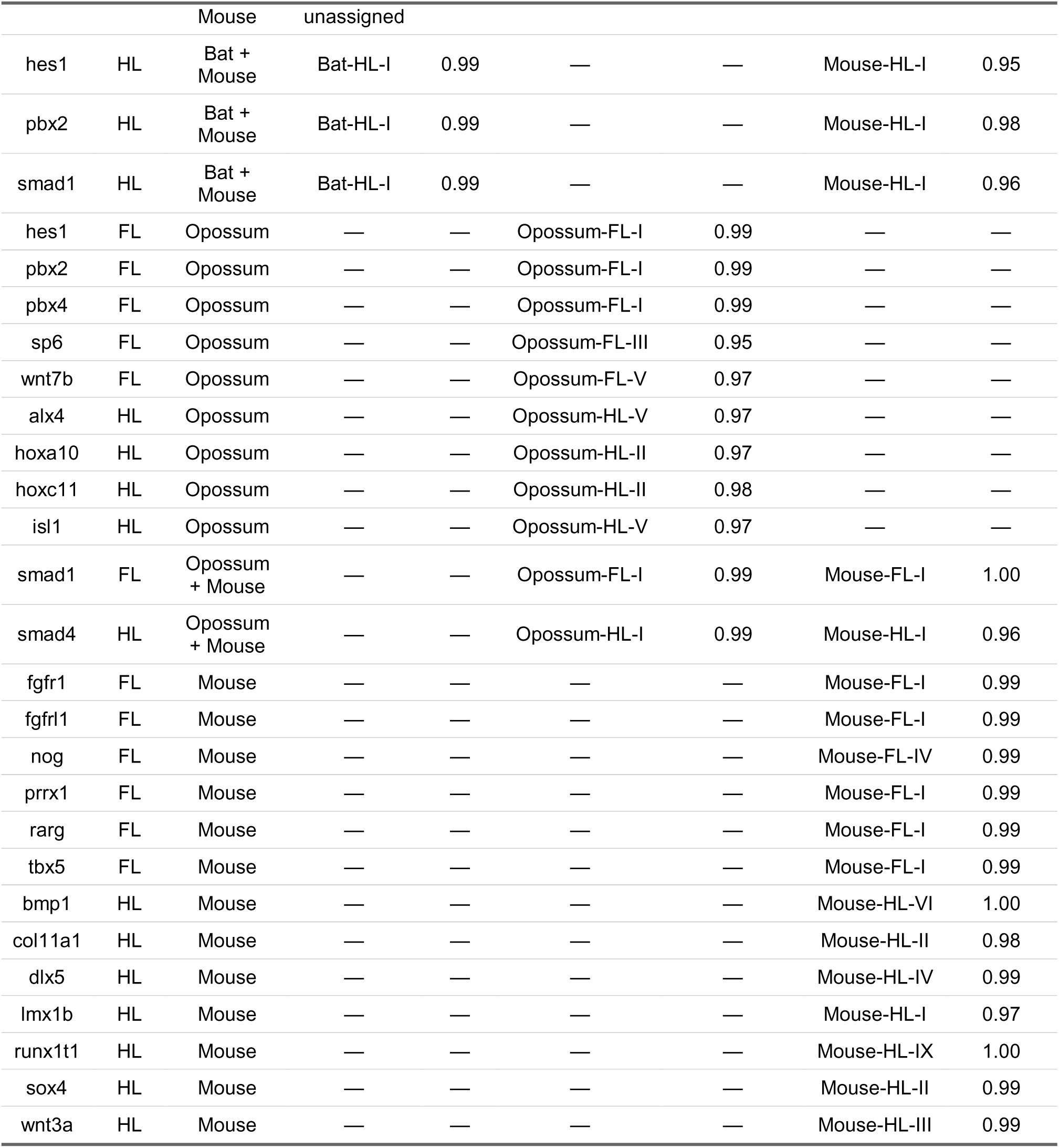
Target limb-development genes identified as hubs in at least one but not all three species. Hub genes defined by exhibiting top 10% kME within their assigned module. Bat- specific hubs constitute the largest set, particularly in the forelimb network; mouse and opossum each contribute smaller sets of species-specific hubs. Dashes indicate that the gene is not a hub in that species.

## Discussion

### Bat limbs, in general, exhibit distinct gene co-expression networks

Bat fore- and hind limb co-expression networks show greater dispersion of limb-development genes across modules than mouse or opossum, and the developmental categories we test show the same pattern (Figures 2 and 3, Table 1). In contrast, mouse and opossum networks show a relatively more uneven distribution of limb-development genes and categories, with most target genes identified in the largest module. However, across all sampled taxa, few specific module-category enrichments are identified, indicating that limb-development genes are widely distributed across modules in bat, mouse, and opossum fore- and hind limbs (Table 2).

Bat co-expression networks also contain 3 times more limb-development genes identified as hubs (i.e., that exhibit high kME, indicating strong correlation with their assigned module), than those of mouse or opossums (Table 4), most of which are associated with digit patterning (5’ Hox genes), skeletal growth (BMPs and associated factors), and AER formation. Bat networks also have 4-10 times more unassigned limb-development genes that do not strongly correlate with any identified module (Table 3). This contrast, together with the broader dispersion of gene categories across bat networks, is consistent with a divergence of gene co-expression patterns within bat limbs relative to those of other examined mammals.

### Bat forelimbs, in particular, exhibit global differences in gene co-expression

Bat forelimbs are highly modified relative to the generalized mammalian condition – the long bones are elongated, the hand plate is extended and digit patterning is reshaped, and the interdigit membranes are retained. Previously, these differences have been proposed to be associated with the rewiring of developmental networks (Hockman et al., 2008; Cooper & Sears, 2013) and changes in regulatory logic (Cretekos et al., 2008; Booker et al., 2016; Eckalbar et al., 2016), but here we identify a global change in gene co-expression across the entire bat forelimb relative to the more generalized forelimbs of mouse and opossum. The retained interdigital wing membrane, absent in the bat hind limb and both limbs of mouse and opossum, represents a plausible candidate contributor to the dispersion we observe, offering one tissue-level hypothesis for the wider co-expression changes reported here. Recent single-cell analyses of bat limbs refine this picture: rather than resulting from suppressed interdigital apoptosis alone, the wing tissue arises substantially from a distinct fibroblast population marked by *Meis2* and *Tbx3* that repurposes a gene program typically restricted to the proximal limb, while interdigital apoptosis itself remains largely conserved between bats and mice (Schindler et al., 2025). This repurposed proximal program, together with the retained interdigital tissue, offers an additional candidate source of the dispersion in forelimb co-expression networks we observe.

Within our co-expression networks, we identify multiple active co-expression modules within the bat forelimb that may contribute to the unique regulatory and morphological changes associated with bat wing development (Figures 3 and 4). Among these, promising avenues for further interrogation include the uniquely high density of limb-development genes in bat forelimb modules II and III and general distribution of chondrogenic genes across the bat forelimb network, potentially reflecting the rewiring of chondrogenic patterning to accommodate elongated digits, another candidate mechanism alongside interdigital tissue retention. This result is supported by single-cell analyses of a different bat (*Rhinolophus sinicus*) extending to later stages of forelimb and hind limb development, which found prolonged chondrogenesis and delayed osteogenesis in the bat forelimb relative to its hind limb and mouse (Lyu et al., 2025). We also identify several candidate genes that may contribute to the altered developmental architecture underlying bat wing formation, including the AER-related gene *Trp63* and the chondrogenic-related genes *Bmp6*, and *Gdf5*, all of which represent promising targets for future functional and regulatory investigation. This contrasts with mouse networks, where the unique hub genes are known master regulators of limb initiation, which highlights the novelty of the hub genes we identify in bats (Table 4). However, our findings suggest that the novel limb phenotype of the bat forelimb is associated not with a few localized changes, but with a largescale reorganization of co-expression across the limb development program that facilitates the generation of a uniquely modified wing. Furthermore, because our results do not reveal a clear partitioning of specific limb-developmental programs among modules, it remains unclear how this modular organization contributes to the observed developmental and phenotypic differences. This uncertainty in part reflects that our analyses are based on gene co-expression alone, the underlying regulatory interactions remain unresolved, leaving the mechanistic basis of these network changes to be investigated *in situ*.

### Opossum limbs exhibit limited differences in gene co-expression

We identify an enrichment for AER development in the opossum forelimb, which is notable given that opossums do not form a morphologically distinct AER during the relatively rapid and precocial development of the forelimb (Doroba & Sears, 2010; Keyte & Smith, 2010, 2014). The significant enrichment of this category within the opossum’s forelimb module-III is particularly striking because so few modules are enriched for specific limb-development categories. However, this enrichment does not indicate a clear timing-associated difference, as might be expected based on the novel developmental strategies of marsupials. Instead, the broadly similar patterns of module dispersion and enrichment observed in opossum and mouse target-gene networks and not in the broader co-expression network suggest that relative heterochrony driving marsupial limb development may arise through processes other than large-scale reorganization of limb developmental gene networks. We speculate this may instead represent changes in the timing of deployment of a more conserved developmental program, reflected by the divergent physiological trajectories of forelimb and hindlimb tissue during this period (Keyte & Smith, 2010, 2014). This is consistent with single-cell evidence from early opossum organogenesis, which found that anterior developmental programs initiate earlier and progress faster than corresponding morphological landmarks, producing an uncoupling of transcriptional and morphological timelines rather than an entire reorganization of developmental gene networks (Menchero et al., 2025).

In contrast to the broad co-expression differences identified in bats, opossum limb morphology does not differ substantially from that of the mouse in the way that bat forelimbs diverge from generalized mammalian morphology. Instead, further study of genes such as *Pbx2* and *Isl1*, which specifically mark forelimb and hind limb initiation and exhibit distinct module correlations, may provide insight into the mechanisms underlying altered developmental timing in marsupials. This pattern suggests that more extensive changes to developmental network organization may be required to generate major morphological innovations, whereas comparatively subtle changes may be sufficient to alter developmental timing.

### Strengths, limitations, and future directions of comparative network analysis

Our work supports a model in which changes in gene co-expression, identifiable through bulk RNA-seq data, may reflect alterations in developmental architecture associated with the evolution of novel phenotypes. These findings highlight WGCNA as a powerful tool for comparative analyses of development across species, while also underscoring several limitations inherent to our approach. Though we address this in our study via the consensus-based network generation that WGCNA performs, sampling depth and breadth limit the power of our analyses, as is common in studies of non-model taxa. Furthermore, our functional categories are mostly informed by work performed on mice, leaving the possibility that genes unique to opossum or bat limb development have been excluded. We did not define categories directly from GO annotations, as broad GO-term membership typically includes genes with indirect or pleiotropic roles that would dilute the specificity of direct cross-species comparison; the narrow, limb-development-specific GO enrichment analysis we do apply (Table S4b) instead serves as an independent validation check on our already-constructed networks, rather than a basis for defining the target gene set itself. While our approach is simplified by focusing on targets with functionally tested roles in limb development, it necessarily excludes important nuances that could emerge from more detailed, species-specific analyses such as recent single-cell studies of bat, opossum, and human limb development (Zhang et al., 2023; Lyu et al., 2025; Menchero et al., 2025; Schindler et al., 2025). These cell-type resolved atlases offer a promising basis for future redefinition of our functional categories, though at some cost to the direct functional validation our curated gene set currently provides. Further, our curated gene set exposes a potential inconsistency in the mouse results that cannot be directly explained. We identify different patterns of modularity between the forelimb and hind limb in the target-gene network that are not reflected in the whole-gene network of the mouse, which is not identified in any of our other taxa. This may reflect a bias introduced by the curated gene set, which is primarily constructed for functional experiments on mice. Broader gene sets, particularly from the referenced, multi-taxon cell-type atlases, or validation of this pattern across other limb sequencing datasets would clarify the source of this unexplained result.

Finally, because WGCNA module assignments are sensitive to data quality and soft-thresholding power, and because the data for this study were generated using bulk RNA-seq on older sequencing platforms, some module-assignment instability may exist that could be refined with modern sequencing approaches. Consistent with this, the mouse forelimb-to-hindlimb redistribution of target genes among modules described above is not mirrored in whole-network module structure and remains without a confirmed causal explanation, though it correlates with a modest, non-definitive shift toward hindlimb-specific chondrogenesis-associated connectivity (Table S5). If this does not reflect bias from our curated gene set, we speculate that differences in the early regulatory architecture of forelimb and hind limb may be contributing to this pattern; a more detailed analysis outside the scope of this work integrating single cell and bulk RNA-seq and ATAC-seq would help resolve this question. Bat gene annotations were derived from a TOGA-based projection rather than a fully de novo *Carollia perspicillata* annotation; while this approach is standard for non-model genomes and none of our limb-development target genes are affected by non-one-to-one orthology classifications in the broader annotated gene set, future work incorporating multi-reference or expression-informed bat annotations may further refine gene recovery and orthology assignment.

Together, our results suggest that divergence in co-expression network architecture may represent an important feature of developmental and phenotypic evolution. Using WGCNA, we identify broad differences in co-expression organization across mammalian limb development, including extensive changes associated with the highly derived bat forelimb and comparatively limited differences associated with marsupial limb heterochrony. These findings support the idea that major morphological innovations may require more extensive alterations to developmental network organization than shifts in developmental timing alone. More broadly, this work highlights the utility of comparative co-expression analyses for investigating developmental evolution in non-model organisms and identifies candidate genes and pathways that provide a foundation for future functional and regulatory studies of mammalian limb diversification.

