## Supplementary Figures and Tables for "Gene co-expression networks reveal differential developmental modularity in Mammalian limbs"

Supplementary Figures S1–S4: Network Dendrograms

Referenced in Methods (Network generation). Branch height reflects dissimilarity (1 − topological overlap); colored bars below each dendrogram indicate final module assignment after merging. Module colors are assigned independently within each network and do not indicate correspondence across networks or figures but correspond to size-rank (e.g. turquoise module is the largest module in each network, blue the next largest). As the species-specific dendrograms presented here are only one block of many, they are not guaranteed to show the largest module dominating the dendrogram, though the ranking holds across the full network. Module color has been renamed to roman numerals, a color-to-numeral correspondence table is not included here, as individual modules are not referenced by name in these supplementary figures.

Figure S1. **Consensus network dendrograms with module color assignments.**

Networks were constructed using blockwiseConsensusModules (Langfelder & Horvath, 2008) with maxBlockSize = 15000, so each dendrogram shows the full network (all genes and module assignments in a single block). Block sizes: Consensus FL — 10,562 genes; Consensus HL — 10,554 genes. WGCNA assigns module color by size (e.g., turquoise is the largest module, blue the second largest, etc.). Because this network is built in a single block, this ranking is fully reflected in the module composition shown here; dendrogram leaf order itself, however, is determined by clustering, not by module size.

**(A)** Forelimb (FL) consensus network.

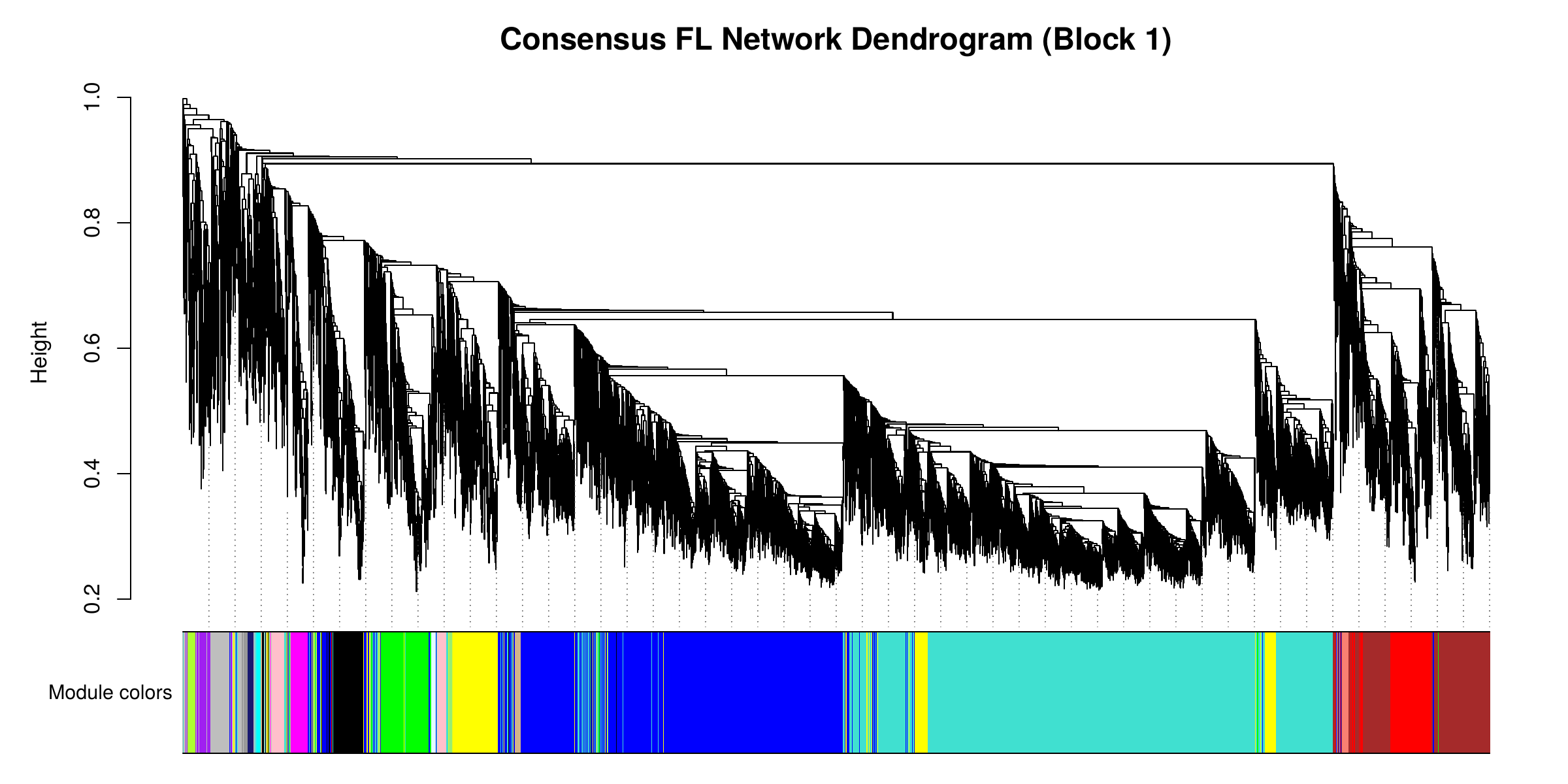

**(B)** Hindlimb (HL) consensus network.

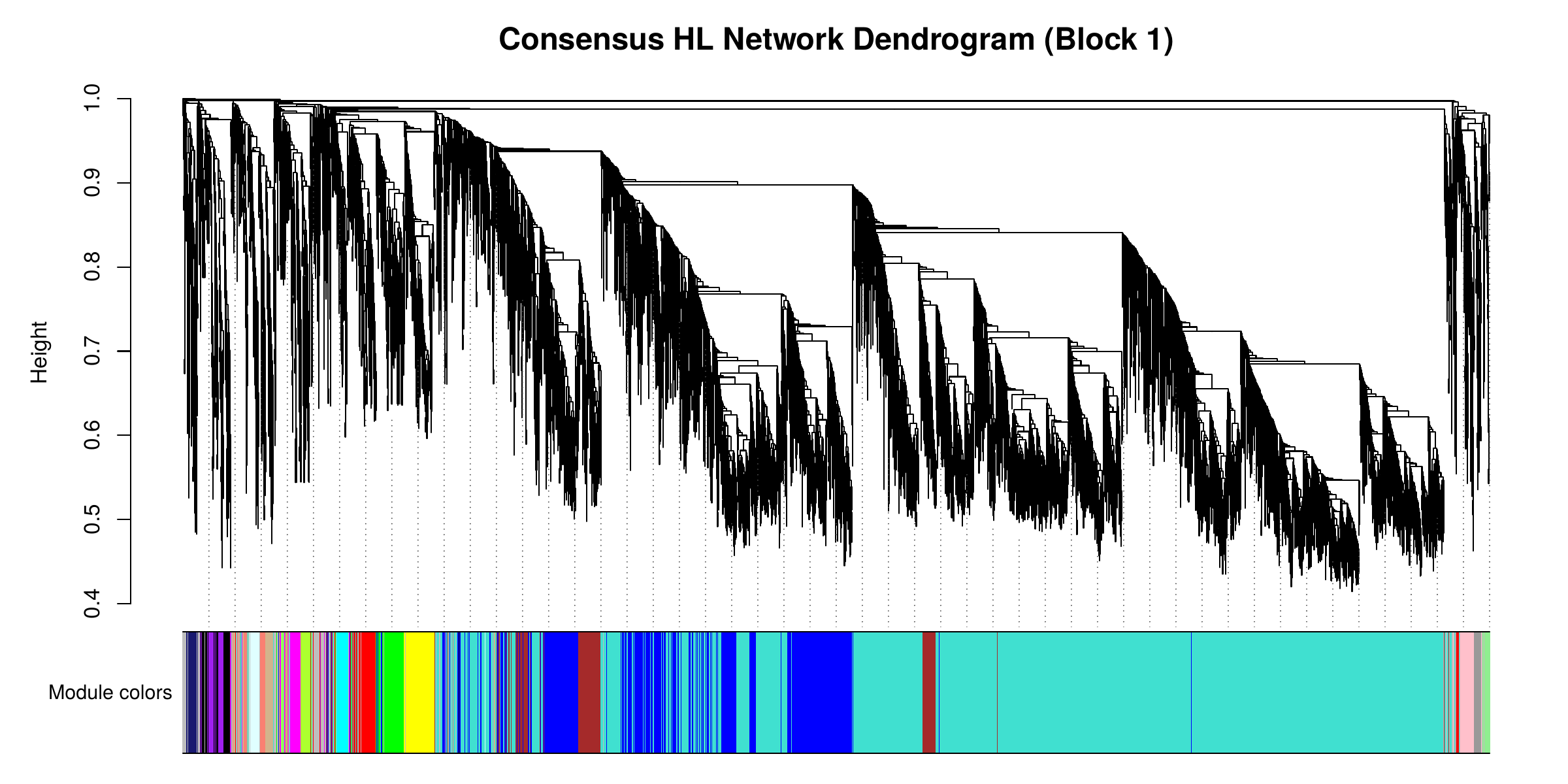

Figure S2. **Bat (Carollia perspicillata) species-specific network dendrograms.**

Networks were constructed using blockwiseModules with the default maxBlockSize = 5000. Each dendrogram shows gene clustering for the largest block (Block 1) only, as the full network was constructed across multiple blocks; the full module structure used in main-text analyses spans all blocks. Block sizes: Bat FL – 4931 / 2983 / 2648; Bat HL – 4987 / 4854 / 713.

**(A)** Forelimb (FL) network.

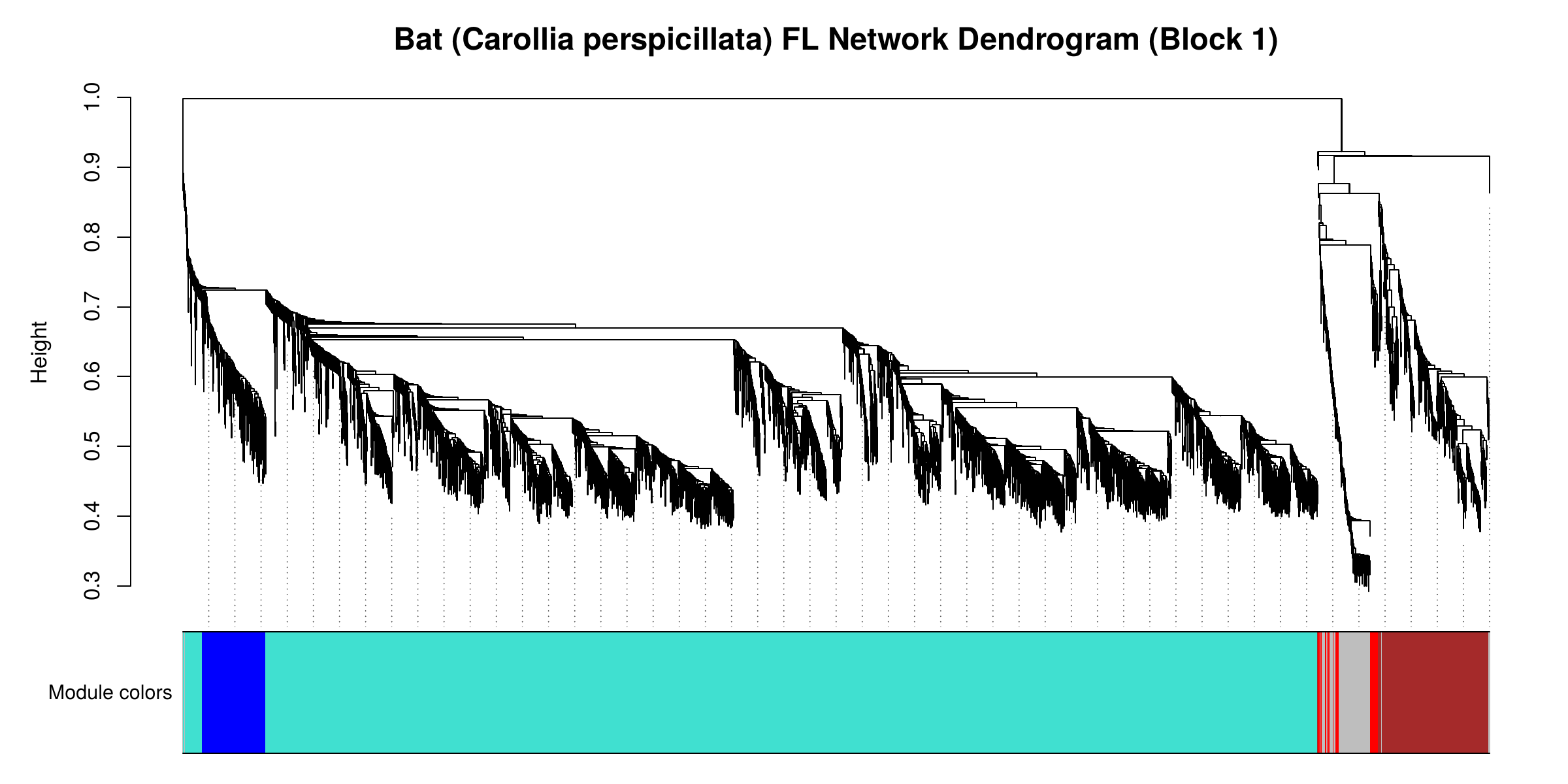

**(B)** Hindlimb (HL) network.

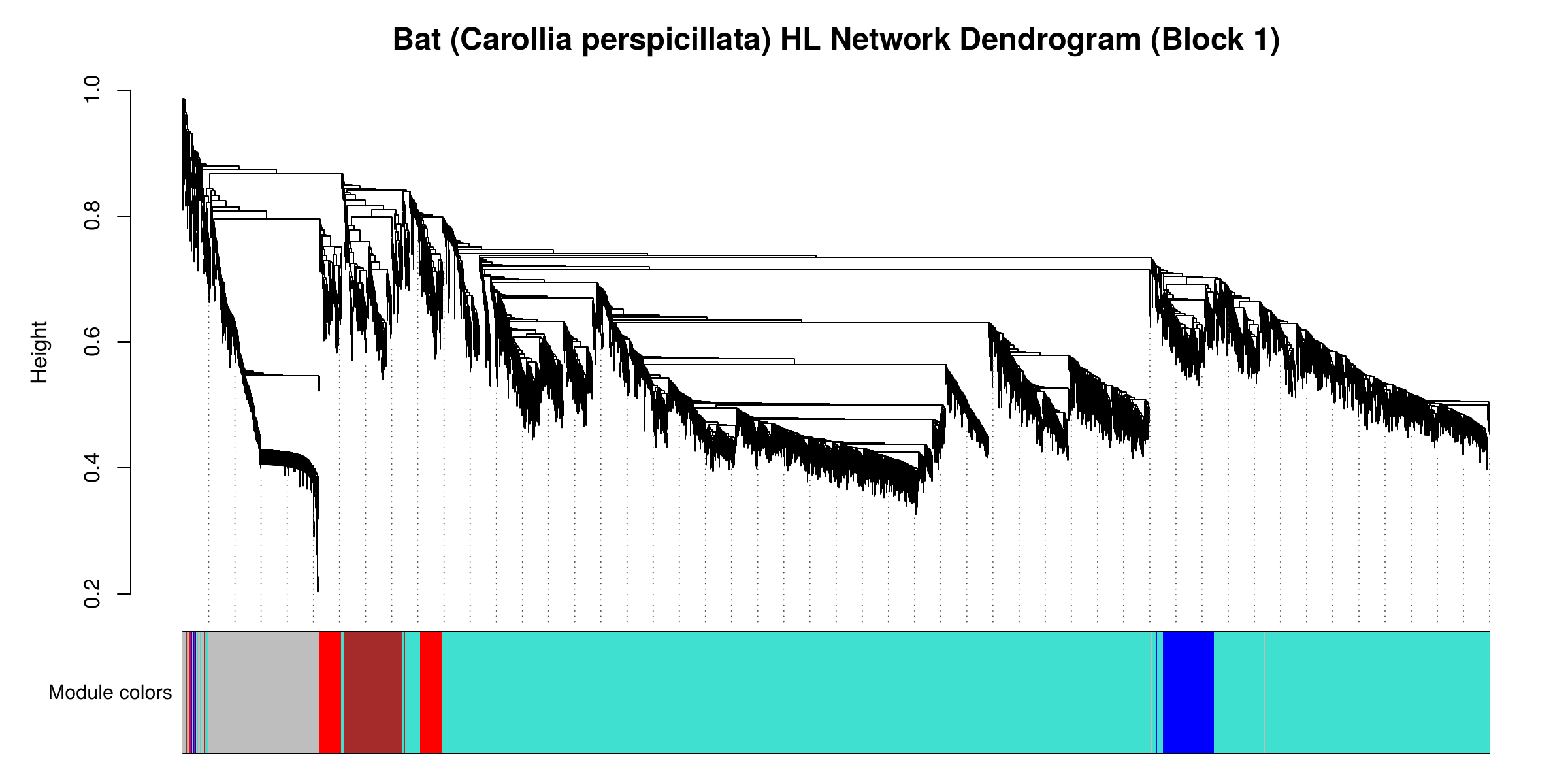

Figure S3. **Opossum (Monodelphis domestica) species-specific network dendrograms.**

Networks were constructed using blockwiseModules with the default maxBlockSize = 5000. Each dendrogram shows gene clustering for the largest block (Block 1) only, as the full network was constructed across multiple blocks; the full module structure used in main-text analyses spans all blocks. Block sizes: Opossum FL – 5000 / 4958 / 604; Opossum HL – 4983 / 4973 / 598.

**(A)** Forelimb (FL) network.

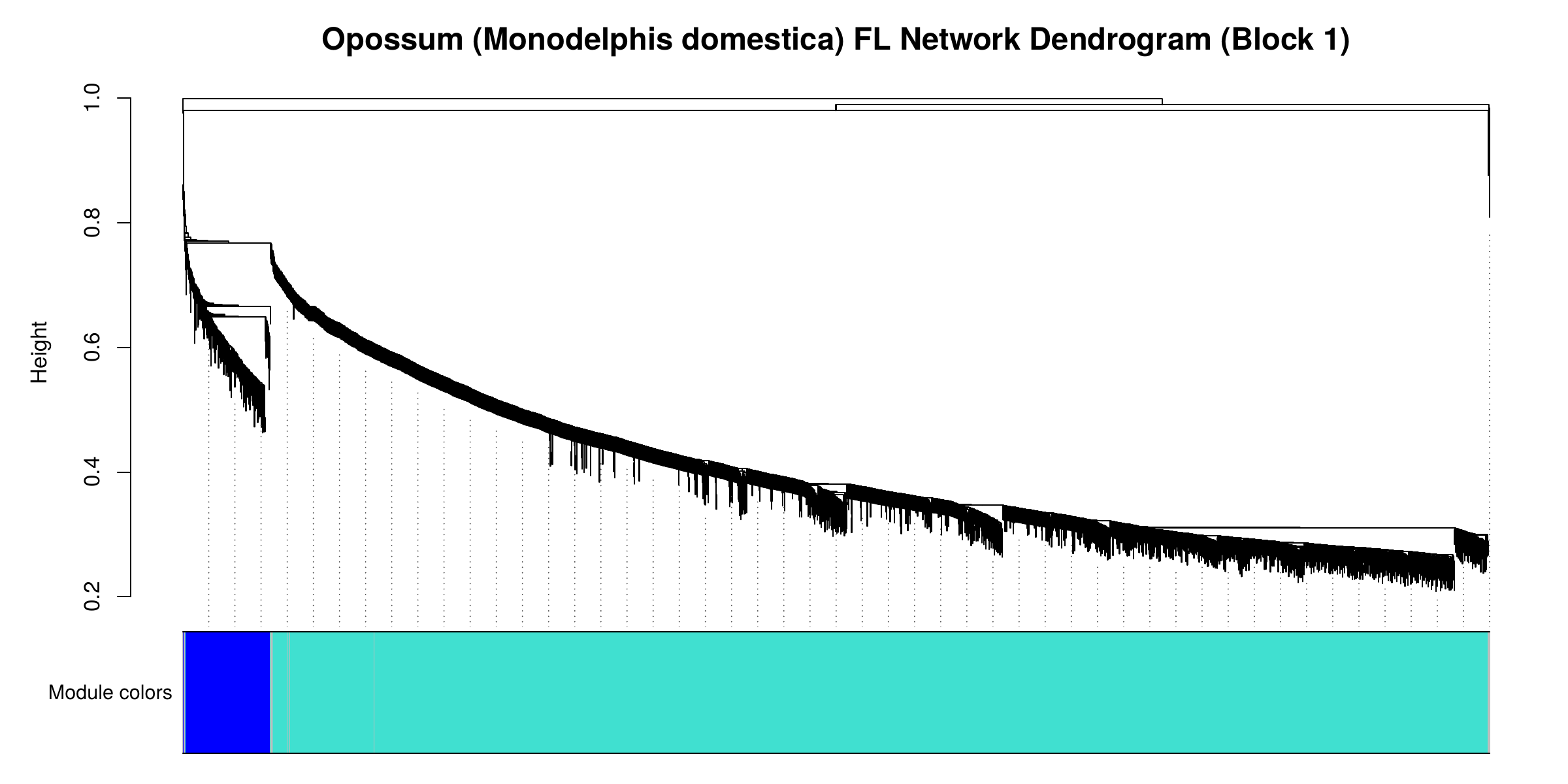

**(B)** Hindlimb (HL) network.

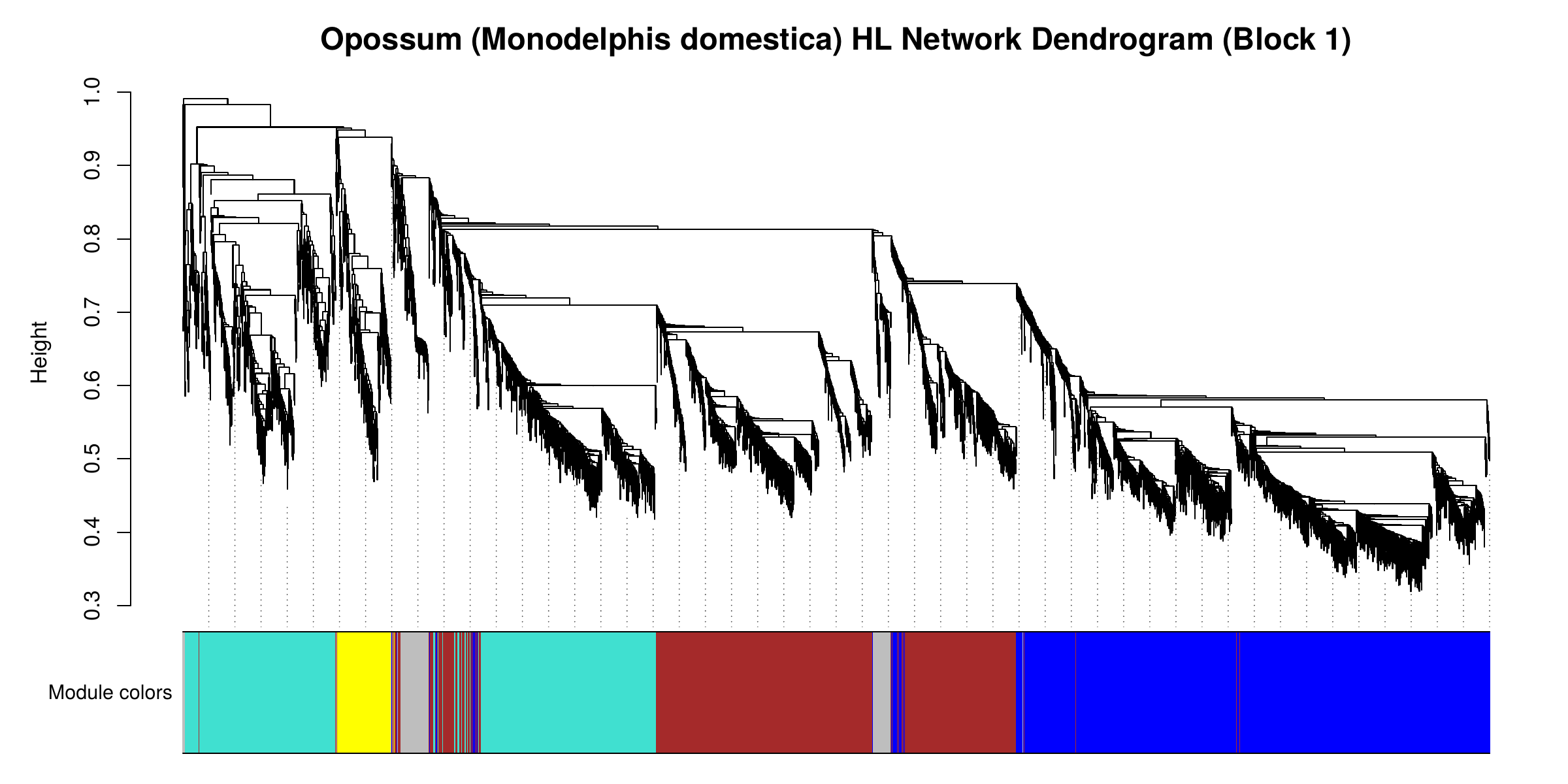

Figure S4. **Mouse (Mus musculus) species-specific network dendrograms.**

Networks were constructed using blockwiseModules with the default maxBlockSize = 5000. Each dendrogram shows gene clustering for the largest block (Block 1) only, as the full network was constructed across multiple blocks; the full module structure used in main-text analyses spans all blocks. Block sizes: Mouse FL – 4965 / 4754 / 843; Mouse HL – 4338 / 4145 / 2071.

**(A)** Forelimb (FL) network.

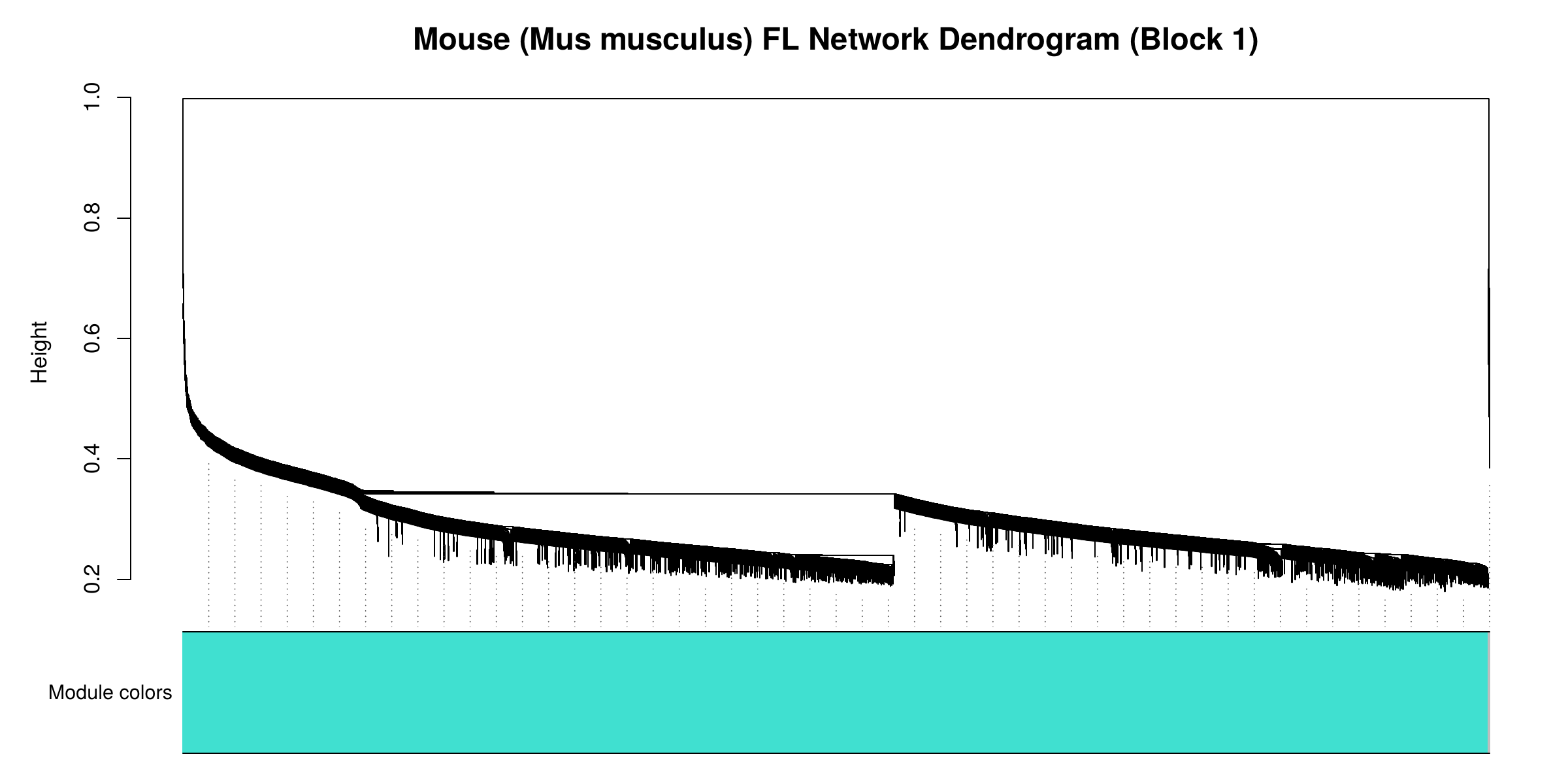

**(B)** Hindlimb (HL) network.

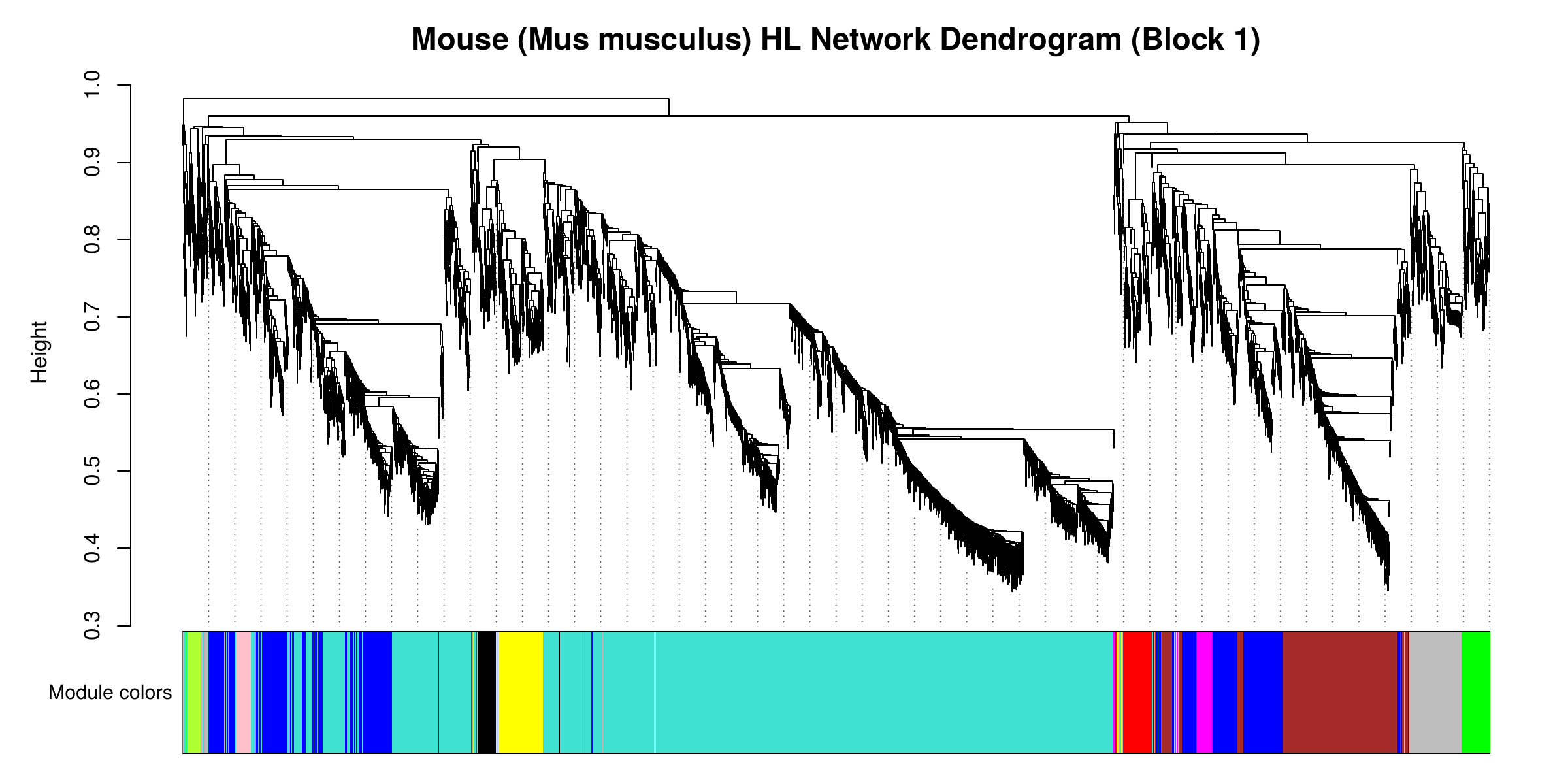

Supplementary Tables S1 – S8a

Comparative WGCNA analysis of limb development in bat, opossum, and mouse. Tables S1–S8a provide gene assignments, network parameters, GO and category enrichment results, and dispersion analyses supporting the main text. See Methods for analytical details.

Table S1. **Curated limb-development gene set used in functional category analyses.** Gene assignments to functional categories were drawn from literature review of limb development across mammals. Some genes are assigned to multiple categories reflecting pleiotropic function in limb development.

| **Gene** | **Category** | **Reference** |
| --- | --- | --- |
| axin2 | AER_signaling | (Jho et al., 2002; Barrow et al., 2003; Ten Berge et al., 2008) |
| bmp2 | AER_signaling | (Zou & Niswander, 1996; Bandyopadhyay et al., 2006; Sears et al., 2006) |
| bmp4 | AER_signaling | (Zou & Niswander, 1996; Bandyopadhyay et al., 2006; Sears et al., 2006; Cooper et al., 2014; Eckalbar et al., 2016) |
| bmp7 | AER_signaling | (Sears et al., 2006; Zeller et al., 2009; Choi et al., 2012) |
| bmpr1a | AER_signaling | (Ovchinnikov et al., 2006; Robert, 2007; Eckalbar et al., 2016) |
| dkk1 | AER_signaling | (Grotewold et al., 1999; Grotewold & Rüther, 2002; Adamska et al., 2004; Díaz-Hernández et al., 2014) |
| dlx2 | AER_signaling | (Kraus & Lufkin, 2006; Conte et al., 2016) |
| dlx3 | AER_signaling | (Sumiyama & Ruddle, 2003) |
| dlx5 | AER_signaling | (Kraus & Lufkin, 2006; Levi et al., 2022) |
| dlx6 | AER_signaling | (Kraus & Lufkin, 2006; Levi et al., 2022) |
| en1 | AER_signaling | (Kimmel et al., 2000; Huettl et al., 2015) |
| etv4 | AER_signaling | (Mao et al., 2009; Zhang et al., 2009) |
| etv5 | AER_signaling | (Mao et al., 2009; Zhang et al., 2009) |
| fgf17 | AER_signaling | (Lewandoski et al., 2000; Ornitz & Itoh, 2015) |
| fgf2 | AER_signaling | (Fallon et al., 1994) |
| fgf20 | AER_signaling | (Hajihosseini & Heath, 2002; Van Greenen & Hockman, 2024) |
| fgf4 | AER_signaling | (G. R. Martin, 1998; Sun et al., 2002; Ornitz & Itoh, 2015) |
| fgf8 | AER_signaling | (Lewandoski et al., 2000; Moon & Capecchi, 2000; Sun et al., 2002; Ornitz & Itoh, 2015) |
| fgfr2 | AER_signaling | (Yu & Ornitz, 2008; Su et al., 2014; Jin et al., 2019) |
| fgfrl1 | AER_signaling | (Wiedemann & Trueb, 2000; Bertrand et al., 2009) |
| grem1 | AER_signaling | (Khokha et al., 2003; Michos et al., 2004; Zuniga, 2015; Eckalbar et al., 2016) |
| hoxc12 | AER_signaling | (Fernandez-Guerrero et al., 2020) |
| hoxc13 | AER_signaling | (Fernandez-Guerrero et al., 2020) |
| hoxc5 | AER_signaling | (Fernandez-Guerrero et al., 2020) |
| hoxc8 | AER_signaling | (Fernandez-Guerrero et al., 2020) |
| isl1 | AER_signaling | (Narkis et al., 2012; Rabinowitz & Vokes, 2012) |
| jag2 | AER_signaling | (Jiang et al., 1998; Pan et al., 2005) |
| msx1 | AER_signaling | (Bensoussan‐Trigano et al., 2011; Markman et al., 2023) |
| msx2 | AER_signaling | (Zeller et al., 2009; Bensoussan‐Trigano et al., 2011) |
| notch1 | AER_signaling | (Francis et al., 2005; Pan et al., 2005) |
| notch2 | AER_signaling | (Pan et al., 2005; Dong et al., 2010) |
| perp | AER_signaling | (Ihrie et al., 2005) |
| sox14 | AER_signaling | (Wilmore et al., 2000) |
| sp6 | AER_signaling | (Hertveldt et al., 2008; Haro et al., 2014) |
| sp8 | AER_signaling | (S. M. Bell et al., 2003; Kawakami et al., 2004; Hertveldt et al., 2008) |
| sp9 | AER_signaling | (Kawakami et al., 2004) |
| tbx3 | AER_signaling | (Washkowitz et al., 2012; Sheeba & Logan, 2017; Soussi et al., 2024) |
| trp63 | AER_signaling | (Mills et al., 1999; Kawata et al., 2017) |
| wnt10a | AER_signaling | (J. Wang & Shackleford, 1996; Witte et al., 2009) |
| wnt10b | AER_signaling | (J. Wang & Shackleford, 1996; Kantaputra et al., 2018) |
| wnt11 | AER_signaling | (Summerhurst et al., 2008) |
| wnt3 | AER_signaling | (Barrow et al., 2003; Glotzer et al., 2022) |
| wnt5b | AER_signaling | (Witte et al., 2009; Vlashi et al., 2023) |
| wnt6 | AER_signaling | (Geetha-Loganathan et al., 2010; Fernandez‐Guerrero et al., 2022) |
| alx4 | AP_patterning | (S. Qu et al., 1997; Kuijper et al., 2005) |
| dkk1 | AP_patterning | (Grotewold et al., 1999; Grotewold & Rüther, 2002; Adamska et al., 2004; Díaz-Hernández et al., 2014) |
| ets1 | AP_patterning | (Lettice et al., 2012) |
| etv4 | AP_patterning | (Mao et al., 2009; Zhang et al., 2009) |
| etv5 | AP_patterning | (Mao et al., 2009; Zhang et al., 2009) |
| gli1 | AP_patterning | (Ahn & Joyner, 2004) |
| gli2 | AP_patterning | (Büscher & Rüther, 1998) |
| gli3 | AP_patterning | (Zeller et al., 2009; Lopez-Rios, 2016) |
| grem1 | AP_patterning | (Michos et al., 2004; Zuniga, 2015) |
| hand1 | AP_patterning | (Barnes et al., 2010; Firulli et al., 2017) |
| hand2 | AP_patterning | (Benazet & Zeller, 2009; Petit et al., 2017) |
| hes1 | AP_patterning | (Bhat et al., 2019; Sharma et al., 2021) |
| hhip | AP_patterning | (Chuang & McMahon, 1999) |
| hoxa10 | AP_patterning | (Favier et al., 1996; Woltering et al., 2014) |
| hoxa13 | AP_patterning | (Fromental-Ramaint et al., 1995) |
| hoxb9 | AP_patterning | (Pineault & Wellik, 2014) |
| hoxd12 | AP_patterning | (Knezevic et al., 1997; Tickle & Towers, 2017) |
| hoxd13 | AP_patterning | (Zeller et al., 2009; Tickle & Towers, 2017) |
| irx3 | AP_patterning | (D. Li et al., 2014) |
| irx5 | AP_patterning | (D. Li et al., 2014) |
| lhx2 | AP_patterning | (Rodriguez-Esteban et al., 1998; Tzchori et al., 2009) |
| meis1 | AP_patterning | (Delgado et al., 2021) |
| pax9 | AP_patterning | (McGlinn et al., 2005) |
| pbx2 | AP_patterning | (Capellini et al., 2006) |
| ptch1 | AP_patterning | (Butterfield et al., 2009) |
| shh | AP_patterning | (Hockman et al., 2008; Benazet & Zeller, 2009; Chew et al., 2014; Tickle & Towers, 2017) |
| smo | AP_patterning | (Long et al., 2001; Varjosalo & Taipale, 2008) |
| tbx2 | AP_patterning | (Farin et al., 2013) |
| tbx3 | AP_patterning | (Washkowitz et al., 2012; Sheeba & Logan, 2017; Soussi et al., 2024) |
| acan | Chondrogenesis | (Lauing et al., 2014; Schwartz & Domowicz, 2022) |
| bmp1 | Chondrogenesis | (Hall, 2015) |
| bmp2 | Chondrogenesis | (Zou & Niswander, 1996; Bandyopadhyay et al., 2006; Sears et al., 2006) |
| bmp3 | Chondrogenesis | (Bahamonde & Lyons, 2001; Gamer et al., 2008; Eckalbar et al., 2016) |
| bmp5 | Chondrogenesis | (Hall, 2015; Eckalbar et al., 2016) |
| bmp6 | Chondrogenesis | (Hall, 2015) |
| bmpr1a | Chondrogenesis | (Ovchinnikov et al., 2006; Robert, 2007; Eckalbar et al., 2016) |
| bmpr1b | Chondrogenesis | (Eckalbar et al., 2016) |
| ccne1 | Chondrogenesis | (Ali-Khan & Hales, 2006) |
| cdk1 | Chondrogenesis | (Saito et al., 2016) |
| cdk6 | Chondrogenesis | (Ito et al., 2014) |
| chrd | Chondrogenesis | (Lorda-Diez et al., 2013; Mouri et al., 2018) |
| col10a1 | Chondrogenesis | (Kwan et al., 1997; Hall, 2015) |
| col11a1 | Chondrogenesis | (Y. Li et al., 1995; Fernandes et al., 2007; Hafez et al., 2015) |
| col2a1 | Chondrogenesis | (S.-W. Li, 1995; D. M. Bell et al., 1997) |
| cyp26b1 | Chondrogenesis | (Pennimpede et al., 2010; Dranse et al., 2011; Probst et al., 2011) |
| dkk1 | Chondrogenesis | (Grotewold et al., 1999; Grotewold & Rüther, 2002; Adamska et al., 2004; Díaz-Hernández et al., 2014) |
| dlx2 | Chondrogenesis | (Levi et al., 2022; S. C. Xu et al., 2001) |
| dlx3 | Chondrogenesis | (Ghoul-Mazgar et al., 2005; Isaac et al., 2014; Levi et al., 2022) |
| dlx5 | Chondrogenesis | (Kraus & Lufkin, 2006; Levi et al., 2022) |
| dlx6 | Chondrogenesis | (Kraus & Lufkin, 2006; Levi et al., 2022) |
| fgf18 | Chondrogenesis | (Liu et al., 2002; Ohbayashi et al., 2002; Hung et al., 2016) |
| fgf9 | Chondrogenesis | (Hung et al., 2007, 2016) |
| fgfr1 | Chondrogenesis | (Su et al., 2014) |
| fgfr2 | Chondrogenesis | (Yu & Ornitz, 2008; Su et al., 2014; Jin et al., 2019) |
| fgfr3 | Chondrogenesis | (Su et al., 2014; Hall, 2015) |
| fgfr4 | Chondrogenesis | (Su et al., 2014) |
| fgfrl1 | Chondrogenesis | (Wiedemann & Trueb, 2000; Bertrand et al., 2009) |
| gdf5 | Chondrogenesis | (Baur et al., 2000; Hall, 2015; Eckalbar et al., 2016) |
| gdf6 | Chondrogenesis | (Settle et al., 2003) |
| gli1 | Chondrogenesis | (Büscher & Rüther, 1998; Ahn & Joyner, 2004) |
| gli2 | Chondrogenesis | (Mo et al., 1997; Büscher & Rüther, 1998) |
| grem1 | Chondrogenesis | (Merino et al., 1999; Lancman et al., 2022) |
| hes1 | Chondrogenesis | (Dong et al., 2010; Rutkowski et al., 2016; Sharma et al., 2021) |
| hoxa11 | Chondrogenesis | (Zeller et al., 2009; Gross et al., 2012) |
| hoxa13 | Chondrogenesis | (Fromental-Ramaint et al., 1995; Zeller et al., 2009) |
| hoxd10 | Chondrogenesis | (Carpenter et al., 1997; Tickle & Towers, 2017) |
| hoxd9 | Chondrogenesis | (Raines et al., 2015; B. Xu & Wellik, 2011) |
| igf1 | Chondrogenesis | (Agrogiannis et al., 2014; Racine & Serrat, 2020) |
| ihh | Chondrogenesis | (St-Jacques et al., 1999; Amano et al., 2015; Wu et al., 2025) |
| mycc | Chondrogenesis | (Zhou et al., 2011) |
| mycn | Chondrogenesis | (Ota et al., 2007; Zhou et al., 2011) |
| nog | Chondrogenesis | (Brunet et al., 1998; Zehentner et al., 2002) |
| notch1 | Chondrogenesis | (Francis et al., 2005; Pan et al., 2005) |
| prrx1 | Chondrogenesis | (J. F. Martin et al., 1995; J. F. Martin & Olson, 2000; Cretekos et al., 2008) |
| prrx2 | Chondrogenesis | (Ten Berge et al., 1998) |
| pth1r | Chondrogenesis | (Lanske et al., 1996; Chung et al., 1998; Zehentner et al., 2002) |
| pthlh | Chondrogenesis | (Lanske et al., 1996; Minina et al., 2002; Hall, 2007) |
| rarg | Chondrogenesis | (Dolle et al., 1989; Galdones & Hales, 2008; Pennimpede et al., 2010) |
| runx2 | Chondrogenesis | (Cobb et al., 2006; Komori, 2011) |
| runx3 | Chondrogenesis | (Soung et al., 2007; Yoshida et al., 2004) |
| smad1 | Chondrogenesis | (Retting et al., 2009) |
| smad4 | Chondrogenesis | (Lim et al., 2015) |
| smad5 | Chondrogenesis | (Retting et al., 2009) |
| smad6 | Chondrogenesis | (Vargesson & Laufer, 2009; Estrada et al., 2011) |
| smad9 | Chondrogenesis | (Retting et al., 2009; Song et al., 2009) |
| smo | Chondrogenesis | (Long et al., 2001; Varjosalo & Taipale, 2008) |
| sox11 | Chondrogenesis | (Bhattaram et al., 2014; Lefebvre & Bhattaram, 2016) |
| sox13 | Chondrogenesis | (Y. Wang et al., 2006) |
| sox4 | Chondrogenesis | (Nissen-Meyer et al., 2007; Bhattaram et al., 2010) |
| sox5 | Chondrogenesis | (Lefebvre, 1998; Smits et al., 2001; Akiyama et al., 2002) |
| sox6 | Chondrogenesis | (Smits et al., 2001; Lefebvre & Smits, 2005; Dy et al., 2010) |
| sox8 | Chondrogenesis | (Molin et al., 2024; González Alvarado & Aprato, 2025) |
| sox9 | Chondrogenesis | (Akiyama et al., 2002, 2007; Molin et al., 2024) |
| tbx15 | Chondrogenesis | (Sheeba & Logan, 2017; Singh et al., 2005) |
| tbx2 | Chondrogenesis | (Farin et al., 2013) |
| wnt16 | Chondrogenesis | (X. Qu et al., 2021; Witte et al., 2009) |
| wnt4 | Chondrogenesis | (Lee & Behringer, 2007) |
| wnt5a | Chondrogenesis | (Gao et al., 2018; Gros et al., 2010) |
| wnt5b | Chondrogenesis | (Vlashi et al., 2023; Witte et al., 2009) |
| wnt9a | Chondrogenesis | (Später et al., 2006) |
| bax | Interdigit_apoptosis | (Bečić et al., 2016; Hernández-Martínez & Covarrubias, 2011) |
| bcl2 | Interdigit_apoptosis | (Bečić et al., 2016; Hernández-Martínez & Covarrubias, 2011) |
| bmp2 | Interdigit_apoptosis | (Bandyopadhyay et al., 2006; Sears et al., 2006; Zou & Niswander, 1996) |
| bmp4 | Interdigit_apoptosis | (Zou & Niswander, 1996; Bandyopadhyay et al., 2006; Sears et al., 2006; Cooper et al., 2014; Eckalbar et al., 2016) |
| bmp7 | Interdigit_apoptosis | (Sears et al., 2006; Zeller et al., 2009; Choi et al., 2012) |
| casp3 | Interdigit_apoptosis | (Kudelova et al., 2012; Kuida et al., 1998) |
| dkk1 | Interdigit_apoptosis | (Grotewold et al., 1999; Grotewold & Rüther, 2002; Adamska et al., 2004; Díaz-Hernández et al., 2014) |
| fgf2 | Interdigit_apoptosis | (Montero et al., 2001) |
| fgf4 | Interdigit_apoptosis | (Lu et al., 2006; Pajni-Underwood et al., 2007; Hernández-Martínez & Covarrubias, 2011) |
| fgf8 | Interdigit_apoptosis | (Pajni-Underwood et al., 2007; Hernández-Martínez & Covarrubias, 2011) |
| gdf10 | Interdigit_apoptosis | (Galdones & Hales, 2008; Gyurján et al., 2011) |
| gdf11 | Interdigit_apoptosis | (Nakashima et al., 1999; Hall, 2015; Matsubara et al., 2017) |
| glis3 | Interdigit_apoptosis | (Kim, 2003) |
| grem1 | Interdigit_apoptosis | (Lancman et al., 2022; Merino et al., 1999) |
| hoxa13 | Interdigit_apoptosis | (Knosp et al., 2004) |
| jag2 | Interdigit_apoptosis | (Jiang et al., 1998; Pan et al., 2005) |
| msx1 | Interdigit_apoptosis | (Bensoussan‐Trigano et al., 2011; Markman et al., 2023) |
| msx2 | Interdigit_apoptosis | (Bensoussan‐Trigano et al., 2011; Zeller et al., 2009) |
| mycn | Interdigit_apoptosis | (Ota et al., 2007) |
| notch1 | Interdigit_apoptosis | (Francis et al., 2005; Pan et al., 2005) |
| notch2 | Interdigit_apoptosis | (Dong et al., 2010; Pan et al., 2005) |
| prrx1 | Interdigit_apoptosis | (Cretekos et al., 2008; J. F. Martin et al., 1995) |
| rdh10 | Interdigit_apoptosis | (Cunningham et al., 2011; Sandell et al., 2007) |
| smad1 | Interdigit_apoptosis | (Wong et al., 2012) |
| smad5 | Interdigit_apoptosis | (Wong et al., 2012) |
| smad6 | Interdigit_apoptosis | (Estrada et al., 2011; Vargesson & Laufer, 2009) |
| tbx2 | Interdigit_apoptosis | (Farin et al., 2013) |
| twist1 | Interdigit_apoptosis | (O’Rourke et al., 2002; Zhang et al., 2010; Loebel et al., 2014; Hirsch et al., 2018) |
| wnt5b | Interdigit_apoptosis | (Vlashi et al., 2023; Witte et al., 2009) |
| aldh1a2 | Limb_initiation | (Mic et al., 2004; Niederreither et al., 2002) |
| dlx1 | Limb_initiation | (Dollé et al., 1992; Hall, 2015) |
| fgf10 | Limb_initiation | (Jin et al., 2019; Min et al., 1998; Sekine et al., 1999) |
| fgfr2 | Limb_initiation | (Yu & Ornitz, 2008; Su et al., 2014; Jin et al., 2019) |
| isl1 | Limb_initiation | (Narkis et al., 2012; Rabinowitz & Vokes, 2012) |
| lhx9 | Limb_initiation | (Yang & Wilson, 2015) |
| meis1 | Limb_initiation | (Delgado et al., 2021; Mercader et al., 2009) |
| meis2 | Limb_initiation | (Capdevila et al., 1999; Mercader et al., 2005; Dai et al., 2014) |
| pbx1 | Limb_initiation | (Capellini et al., 2006, 2008, 2011) |
| pbx2 | Limb_initiation | (Capellini et al., 2006) |
| pitx1 | Limb_initiation | (Logan & Tabin, 1999; Marcil et al., 2003; Nemec et al., 2017) |
| pitx2 | Limb_initiation | (Marcil et al., 2003) |
| rdh10 | Limb_initiation | (Cunningham et al., 2011; Sandell et al., 2007) |
| tbx4 | Limb_initiation | (Sheeba & Logan, 2017; Duboc et al., 2021) |
| tbx5 | Limb_initiation | (Agarwal et al., 2003; Sheeba & Logan, 2017) |
| wnt3a | Limb_initiation | (Kawakami et al., 2001; Lin & Zhang, 2020) |
| wnt8c | Limb_initiation | (Kawakami et al., 2001) |
| bmpr2 | Outgrowth_elongation | (Eckalbar et al., 2016; Gamer et al., 2011) |
| ccnd1 | Outgrowth_elongation | (Ali-Khan & Hales, 2006) |
| etv1 | Outgrowth_elongation | (Yamamoto-Shiraishi et al., 2014) |
| etv4 | Outgrowth_elongation | (Mao et al., 2009; Zhang et al., 2009) |
| etv5 | Outgrowth_elongation | (Mao et al., 2009; Zhang et al., 2009) |
| fgf10 | Outgrowth_elongation | (Jin et al., 2019; Ohuchi et al., 1997) |
| fgf16 | Outgrowth_elongation | (Jamsheer et al., 2013; Rigueur, 2024) |
| fgf2 | Outgrowth_elongation | (Dono & Zeller, 1994; Fallon et al., 1994) |
| fgf9 | Outgrowth_elongation | (Hung et al., 2007, 2016) |
| fgfr1 | Outgrowth_elongation | (Verheyden et al., 2005; X. Xu et al., 1998) |
| gdf5 | Outgrowth_elongation | (Baur et al., 2000; Hall, 2015; Eckalbar et al., 2016) |
| grem1 | Outgrowth_elongation | (Benazet & Zeller, 2009; Merino et al., 1999; Michos et al., 2004) |
| hand2 | Outgrowth_elongation | (Benazet & Zeller, 2009; Petit et al., 2017) |
| hoxd11 | Outgrowth_elongation | (Vargas & Fallon, 2005; Zeller et al., 2009; Tickle & Towers, 2017) |
| igf1 | Outgrowth_elongation | (Agrogiannis et al., 2014; Racine & Serrat, 2020) |
| lhx2 | Outgrowth_elongation | (Rodriguez-Esteban et al., 1998; Tzchori et al., 2009) |
| mycn | Outgrowth_elongation | (Ota et al., 2007; Zhou et al., 2011) |
| prrx1 | Outgrowth_elongation | (Cretekos et al., 2008; J. F. Martin et al., 1995) |
| prrx2 | Outgrowth_elongation | (Ten Berge et al., 1998) |
| runx2 | Outgrowth_elongation | (Cobb et al., 2006; Komori, 2011) |
| shh | Outgrowth_elongation | (Hockman et al., 2008; Benazet & Zeller, 2009; Chew et al., 2014; Tickle & Towers, 2017) |
| tbx4 | Outgrowth_elongation | (Sheeba & Logan, 2017; Duboc et al., 2021) |
| twist1 | Outgrowth_elongation | (O’Rourke et al., 2002; Zhang et al., 2010; Loebel et al., 2014; Hirsch et al., 2018) |
| twist2 | Outgrowth_elongation | (O’Rourke et al., 2002; Wade et al., 2012) |
| wnt16 | Outgrowth_elongation | (X. Qu et al., 2021; Witte et al., 2009) |
| wnt5a | Outgrowth_elongation | (Gao et al., 2018; Gros et al., 2010) |
| cyp26b1 | PD_patterning | (Probst et al., 2011; Yashiro et al., 2004) |
| fgf8 | PD_patterning | (Moon & Capecchi, 2000; Sun et al., 2002) |
| fgf9 | PD_patterning | (Hung et al., 2007, 2016) |
| hand1 | PD_patterning | (Barnes et al., 2010; Firulli et al., 2017) |
| hoxa10 | PD_patterning | (Favier et al., 1996; Woltering et al., 2014) |
| hoxa11 | PD_patterning | (Davis et al., 1995; Wellik & Capecchi, 2003) |
| hoxa13 | PD_patterning | (Fromental-Ramaint et al., 1995) |
| hoxa9 | PD_patterning | (B. Xu & Wellik, 2011) |
| hoxc10 | PD_patterning | (Chen et al., 2024) |
| hoxc11 | PD_patterning | (Papenbrock et al., 2000; Wellik & Capecchi, 2003) |
| hoxc9 | PD_patterning | (Chen et al., 2024) |
| hoxd10 | PD_patterning | (Carpenter et al., 1997; Tickle & Towers, 2017) |
| hoxd11 | PD_patterning | (Zeller et al., 2009; Tickle & Towers, 2017) |
| hoxd13 | PD_patterning | (Zeller et al., 2009; Tickle & Towers, 2017) |
| hoxd9 | PD_patterning | (B. Xu & Wellik, 2011; Raines et al., 2015) |
| meis1 | PD_patterning | (Mercader et al., 2009; Delgado et al., 2021) |
| meis2 | PD_patterning | (Capdevila et al., 1999; Mercader et al., 2005; Dai et al., 2014) |
| pbx1 | PD_patterning | (Capellini et al., 2006, 2008, 2011) |
| pbx2 | PD_patterning | (Capellini et al., 2006) |
| pbx4 | PD_patterning | (Ning et al., 2024) |
| rdh10 | PD_patterning | (Sandell et al., 2007; Cunningham et al., 2011) |
| runx1t1 | PD_patterning | (Yamamoto et al., 2019; Murray et al., 2024) |
| wnt5a | PD_patterning | (Gros et al., 2010; Gao et al., 2018) |

Table S1a. TOGA orthology classification and mouse-homolog mapping for all 18,773 annotated *Carollia perspicillata* genes. Orthology class assigned by TOGA (Kirilenko et al., 2023) using the human (hg38) reference; mouse homolog assigned independently via Ensembl BioMart regardless of TOGA orthology class. None of the 165 curated limb-development target genes (Table S1) are affected by non-one-to-one orthology classification.

Table provided as Supplementary_Table_S1a_Carollia_orthology.csv

Table S2. **Sample composition for comparative WGCNA analysis after quality control filtering.** Expression data were obtained from GEO accession GSE71390. Stage matching performed using supplementary data from Maier et al., 2017. Replicate numbers assigned for this manuscript.

| **Sample ID** | **Species** | **Limb** | **Developmental stage** | **Replicate** | **GEO accession** |
| --- | --- | --- | --- | --- | --- |
| bat_FL_13_rep2 | Carollia perspicillata | FL | Ridge | 1 | GSM2718691 |
| bat_FL_13_rep3 | Carollia perspicillata | FL | Ridge | 2 | GSM2718692 |
| bat_FL_14_rep3 | Carollia perspicillata | FL | Bud | 1 | GSM2718693 |
| bat_FL_ES_rep1 | Carollia perspicillata | FL | Bud | 2 | GSM1833580 |
| bat_FL_ES_rep2 | Carollia perspicillata | FL | Bud | 3 | GSM1833581 |
| bat_FL_LS_rep1 | Carollia perspicillata | FL | Paddle | 1 | GSM1833582 |
| bat_FL_LS_rep2 | Carollia perspicillata | FL | Paddle | 2 | GSM1833583 |
| bat_HL_13_rep2 | Carollia perspicillata | HL | Ridge | 1 | GSM2718695 |
| bat_HL_13_rep3 | Carollia perspicillata | HL | Ridge | 2 | GSM2718696 |
| bat_HL_14_rep1 | Carollia perspicillata | HL | Bud | 1 | GSM2718697 |
| bat_HL_14_rep2 | Carollia perspicillata | HL | Bud | 2 | GSM2718698 |
| bat_HL_14_rep3 | Carollia perspicillata | HL | Bud | 3 | GSM2718699 |
| bat_HL_15_rep1 | Carollia perspicillata | HL | Paddle | 1 | GSM2718700 |
| bat_HL_15_rep2 | Carollia perspicillata | HL | Paddle | 2 | GSM2718701 |
| opossum_FL_27_1 | Monodelphis domestica | FL | Ridge | 1 | GSM2718730 |
| opossum_FL_27_2 | Monodelphis domestica | FL | Ridge | 2 | GSM2718731 |
| opossum_FL_27_3 | Monodelphis domestica | FL | Ridge | 3 | GSM2718732 |
| opossum_FL_ES_rep1 | Monodelphis domestica | FL | Bud | 1 | GSM1833589 |
| opossum_FL_ES_rep2 | Monodelphis domestica | FL | Bud | 2 | GSM1833590 |
| opossum_FL_ES_rep3 | Monodelphis domestica | FL | Bud | 3 | GSM1833591 |
| opossum_FL_ES_rep4 | Monodelphis domestica | FL | Bud | 4 | GSM1833592 |
| opossum_FL_ES_rep5 | Monodelphis domestica | FL | Bud | 5 | GSM1833593 |
| opossum_FL_LS_rep1 | Monodelphis domestica | FL | Paddle | 1 | GSM1833595 |
| opossum_FL_LS_rep2 | Monodelphis domestica | FL | Paddle | 2 | GSM1833596 |
| opossum_FL_LS_rep3 | Monodelphis domestica | FL | Paddle | 3 | GSM1833597 |
| opossum_HL_30_2 | Monodelphis domestica | HL | Ridge | 1 | GSM2718734 |
| opossum_HL_30_3 | Monodelphis domestica | HL | Ridge | 2 | GSM2718735 |
| opossum_HL_31_2 | Monodelphis domestica | HL | Bud | 1 | GSM2718737 |
| opossum_HL_31_3 | Monodelphis domestica | HL | Bud | 2 | GSM2718738 |
| opossum_HL_31_4 | Monodelphis domestica | HL | Bud | 3 | GSM2718739 |
| opossum_HL_32_1 | Monodelphis domestica | HL | Paddle | 1 | GSM2718741 |
| opossum_HL_32_2 | Monodelphis domestica | HL | Paddle | 2 | GSM2718742 |
| mouse_FL_W2_rep2 | Mus musculus | FL | Ridge | 1 | GSM2718703 |
| mouse_FL_W2_rep3 | Mus musculus | FL | Ridge | 2 | GSM2718704 |
| mouse_FL_ES_rep1 | Mus musculus | FL | Bud | 1 | GSM1833584 |
| mouse_FL_ES_rep2 | Mus musculus | FL | Bud | 2 | GSM1833585 |
| mouse_FL_LS_rep1 | Mus musculus | FL | Paddle | 1 | GSM1833586 |
| mouse_FL_LS_rep2 | Mus musculus | FL | Paddle | 2 | GSM1833587 |
| mouse_FL_LS_rep3 | Mus musculus | FL | Paddle | 3 | GSM1833588 |
| mouse_HL_W2_rep1 | Mus musculus | HL | Ridge | 1 | GSM2718706 |
| mouse_HL_W2_rep2 | Mus musculus | HL | Ridge | 2 | GSM2718707 |
| mouse_HL_W3_4_rep1 | Mus musculus | HL | Bud | 1 | GSM2718708 |
| mouse_HL_W3_4_rep2 | Mus musculus | HL | Bud | 2 | GSM2718709 |
| mouse_HL_W3_4_rep3 | Mus musculus | HL | Bud | 3 | GSM2718710 |
| mouse_HL_W6_rep1 | Mus musculus | HL | Paddle | 1 | GSM2718711 |
| mouse_HL_W6_rep2 | Mus musculus | HL | Paddle | 2 | GSM2718712 |
| mouse_HL_W6_rep3 | Mus musculus | HL | Paddle | 3 | GSM2718713 |
| mouse_HL_W6_rep4 | Mus musculus | HL | Paddle | 4 | GSM2718714 |

Table S3. **WGCNA parameters used for network construction.** Consensus networks were constructed using blockwiseConsensusModules() with default correlation settings; species-specific networks were constructed using blockwiseModules() with bicor correlation and outlier tolerance for added robustness (Langfelder & Horvath, 2012). *Consensus networks did not specify corType or maxPOutliers, leaving these at WGCNA defaults (Pearson correlation, no outlier handling).

| **Network** | **Soft power** | **Network type** | **TOM type** | **Min module size** | **Merge cut height** | **Deep split** | **Correlation** | **Max P outliers** |
| --- | --- | --- | --- | --- | --- | --- | --- | --- |
| Consensus FL | 12 | signed | signed | 40 | 0.3 | 2 | Pearson* | default* |
| Consensus HL | 12 | signed | signed | 40 | 0.3 | 2 | Pearson* | default* |
| Bat (Cper) FL | 12 | signed | signed | 40 | 0.3 | 2 | bicor | 0.10 |
| Bat (Cper) HL | 12 | signed | signed | 40 | 0.3 | 2 | bicor | 0.10 |
| Opossum (Mdom) FL | 12 | signed | signed | 40 | 0.3 | 2 | bicor | 0.10 |
| Opossum (Mdom) HL | 12 | signed | signed | 40 | 0.3 | 2 | bicor | 0.10 |
| Mouse (Mmus) FL | 12 | signed | signed | 40 | 0.3 | 2 | bicor | 0.10 |
| Mouse (Mmus) HL | 12 | signed | signed | 40 | 0.3 | 2 | bicor | 0.10 |

Table S4a. **Standard GO Biological Process enrichment, top 5 terms per module per network.** Enrichment computed via clusterProfiler against the org.Mm.eg.db annotation database using mouse-homolog gene symbols. Adjusted p-values use the Benjamini-Hochberg method.

Table provided as Supplementary_Table_S4a_GO_enrichmenttop5_all_networks.csv

Table S4b. **Standard GO Biological Process enrichment filtered to set of limb-development-related GO terms.** Filtering applied post-enrichment to focus on limb-specific signal.

Table provided as Supplementary_Table_S4b_GO_enrichment_limb_all_networks.csv

Table S5. **Fisher's exact test enrichment for all module-category combinations across networks.** Tests are one-sided with each network's expressed-gene set as background. The 'Significant' column flags combinations meeting the strict threshold (overlap >= 5, fold >= 2, within-network adjusted p < 0.05). Adjusted p-values are reported under both within-network FDR (primary) and global FDR across all networks. This table includes the Consensus network in addition to the six species-specific networks summarized in the main-text Table 2; the main-text count of 4 significant enrichments is scoped to the species-specific networks only (Table 1) and does not include the Consensus network result flagged as significant below.

Table provided as Supplementary_Table_S5_module_category_enrichment_all_networks.csv

Table S6. **Per-category dispersion across modules in each network.** Effective N modules computed as exp(Shannon entropy of category gene counts across assigned modules). Evenness ranges 0-1 with 1 indicating perfect spread across occupied modules.

| **Category** | **Genes total** | **Genes assigned** | **Modules occupied** | **Effective N modules** | **Evenness** | **Species** | **Limb** |
| --- | --- | --- | --- | --- | --- | --- | --- |
| AER_signaling | 37 | 29 | 6 | 3.74 | 0.74 | Bat | FL |
| AP_patterning | 26 | 22 | 6 | 4.62 | 0.85 | Bat | FL |
| Chondrogenesis | 60 | 51 | 9 | 6.61 | 0.86 | Bat | FL |
| Interdigit_apoptosis | 25 | 20 | 5 | 3.69 | 0.81 | Bat | FL |
| Limb_initiation | 10 | 8 | 5 | 4.46 | 0.93 | Bat | FL |
| Outgrowth_elongation | 21 | 18 | 7 | 5.61 | 0.89 | Bat | FL |
| PD_patterning | 19 | 18 | 7 | 6.04 | 0.92 | Bat | FL |
| AER_signaling | 38 | 33 | 6 | 4.31 | 0.82 | Bat | HL |
| AP_patterning | 25 | 21 | 5 | 3.48 | 0.78 | Bat | HL |
| Chondrogenesis | 58 | 56 | 8 | 4.18 | 0.69 | Bat | HL |
| Interdigit_apoptosis | 26 | 23 | 5 | 2.51 | 0.57 | Bat | HL |
| Limb_initiation | 10 | 7 | 4 | 3.59 | 0.92 | Bat | HL |
| Outgrowth_elongation | 21 | 18 | 6 | 3.68 | 0.73 | Bat | HL |
| PD_patterning | 19 | 15 | 4 | 2.94 | 0.78 | Bat | HL |
| AER_signaling | 37 | 36 | 4 | 1.59 | 0.33 | Mouse | FL |
| AP_patterning | 26 | 24 | 5 | 1.98 | 0.42 | Mouse | FL |
| Chondrogenesis | 60 | 60 | 4 | 1.69 | 0.38 | Mouse | FL |
| Interdigit_apoptosis | 25 | 25 | 2 | 1.32 | 0.40 | Mouse | FL |
| Limb_initiation | 10 | 10 | 3 | 1.89 | 0.58 | Mouse | FL |
| Outgrowth_elongation | 21 | 19 | 3 | 1.71 | 0.49 | Mouse | FL |
| PD_patterning | 19 | 19 | 3 | 1.51 | 0.37 | Mouse | FL |
| AER_signaling | 38 | 38 | 4 | 1.87 | 0.45 | Mouse | HL |
| AP_patterning | 25 | 23 | 4 | 2.68 | 0.71 | Mouse | HL |
| Chondrogenesis | 58 | 57 | 7 | 3.47 | 0.64 | Mouse | HL |
| Interdigit_apoptosis | 26 | 25 | 5 | 2.80 | 0.64 | Mouse | HL |
| Limb_initiation | 10 | 10 | 3 | 1.89 | 0.58 | Mouse | HL |
| Outgrowth_elongation | 21 | 20 | 4 | 3.03 | 0.80 | Mouse | HL |
| PD_patterning | 19 | 19 | 4 | 2.57 | 0.68 | Mouse | HL |
| AER_signaling | 37 | 37 | 5 | 2.60 | 0.59 | Opossum | FL |
| AP_patterning | 26 | 26 | 4 | 1.62 | 0.35 | Opossum | FL |
| Chondrogenesis | 60 | 60 | 4 | 1.44 | 0.26 | Opossum | FL |
| Interdigit_apoptosis | 25 | 25 | 3 | 1.70 | 0.48 | Opossum | FL |
| Limb_initiation | 10 | 10 | 3 | 1.89 | 0.58 | Opossum | FL |
| Outgrowth_elongation | 21 | 21 | 2 | 1.21 | 0.28 | Opossum | FL |
| PD_patterning | 19 | 19 | 4 | 1.84 | 0.44 | Opossum | FL |
| AER_signaling | 38 | 37 | 6 | 3.68 | 0.73 | Opossum | HL |
| AP_patterning | 25 | 24 | 3 | 2.51 | 0.84 | Opossum | HL |
| Chondrogenesis | 58 | 57 | 5 | 3.87 | 0.84 | Opossum | HL |
| Interdigit_apoptosis | 26 | 25 | 3 | 2.75 | 0.92 | Opossum | HL |
| Limb_initiation | 10 | 10 | 4 | 2.97 | 0.79 | Opossum | HL |
| Outgrowth_elongation | 21 | 20 | 4 | 2.77 | 0.73 | Opossum | HL |
| PD_patterning | 19 | 19 | 3 | 2.36 | 0.78 | Opossum | HL |

Table S7. **Effective number of modules occupied by each category in each network.** Higher values indicate broader distribution. Bat networks show consistently higher dispersion than mouse and opossum networks across most categories.

| **Category** | **Bat_FL** | **Bat_HL** | **Opossum_FL** | **Opossum_HL** | **Mouse_FL** | **Mouse_HL** |
| --- | --- | --- | --- | --- | --- | --- |
| AER_signaling | 3.74 | 4.31 | 2.60 | 3.68 | 1.59 | 1.87 |
| AP_patterning | 4.62 | 3.48 | 1.62 | 2.51 | 1.98 | 2.68 |
| Chondrogenesis | 6.61 | 4.18 | 1.44 | 3.87 | 1.69 | 3.47 |
| Interdigit_apoptosis | 3.69 | 2.51 | 1.70 | 2.75 | 1.32 | 2.80 |
| Limb_initiation | 4.46 | 3.59 | 1.89 | 2.97 | 1.89 | 1.89 |
| Outgrowth_elongation | 5.61 | 3.68 | 1.21 | 2.77 | 1.71 | 3.03 |
| PD_patterning | 6.04 | 2.94 | 1.84 | 2.36 | 1.51 | 2.57 |
| **Average** | 4.97 | 3.53 | 1.76 | 2.99 | 1.67 | 2.62 |

Table S8. **Wilcoxon signed-rank tests for pairwise differences in category dispersion (effective N modules) across networks.** Tests are paired by category (n = 7). Adjusted p-values use the Benjamini-Hochberg method across all 9 comparisons. Note that the minimum achievable p-value is approximately 0.0078 with paired Wilcoxon at n = 7, limiting test resolution.

| **Comparison** | **N categories** | **Median diff** | **Raw p** | **Adj. p (BH)** |
| --- | --- | --- | --- | --- |
| Bat FL vs Mouse FL | 7 | 2.65 | 0.016 | 0.046 |
| Bat FL vs Opossum FL | 7 | 3.00 | 0.016 | 0.046 |
| Opossum FL vs Mouse FL | 7 | 0.00 | 0.834 | 0.834 |
| Bat HL vs Mouse HL | 7 | 0.72 | 0.031 | 0.046 |
| Bat HL vs Opossum HL | 7 | 0.61 | 0.031 | 0.046 |
| Opossum HL vs Mouse HL | 7 | -0.04 | 0.578 | 0.650 |
| Bat FL vs Bat HL | 7 | 1.18 | 0.031 | 0.046 |
| Opossum FL vs Opossum HL | 7 | -1.08 | 0.016 | 0.046 |
| Mouse FL vs Mouse HL | 7 | -1.06 | 0.036 | 0.046 |

Table S8a. **Whole-network module inventory across all eight forelimb and hindlimb co-expression networks (Consensus, Mouse, Opossum, Bat), excluding the unassigned genes.** Reports module count, module size distribution, and largest-module percentage for each network, for direct comparison against the target-gene-level dispersion analyses (Tables S6–S8).

| **Network** | **Species** | **Limb** | **n modules** | **Largest module size** | **Largest module percent** | **Total assigned genes** |
| --- | --- | --- | --- | --- | --- | --- |
| Bat_FL | Bat | FL | 16 | 4310 | 50 | 8627 |
| Mouse_FL | Mouse | FL | 10 | 8279 | 80.8 | 10249 |
| Opossum_FL | Opossum | FL | 5 | 9307 | 90.6 | 10277 |
| Consensus_FL | Consensus | FL | 17 | 3907 | 38.1 | 10259 |
| Bat_HL | Bat | HL | 11 | 5577 | 63.8 | 8741 |
| Mouse_HL | Mouse | HL | 12 | 8187 | 81.1 | 10094 |
| Opossum_HL | Opossum | HL | 9 | 4225 | 41.9 | 10092 |
| Consensus_HL | Consensus | HL | 18 | 6149 | 60.1 | 10236 |
